# Genomic quantification of inbreeding depression in wild vertebrate populations

**DOI:** 10.64898/2026.07.16.737952

**Authors:** Kiran G. L. Lee, Hannah Parkes, Sophie Wilkins, Helen Hipperson, Terry Burke

## Abstract

Inbreeding depression is a critical driver of extinction risk in fragmented populations, yet its magnitude in the wild has historically been estimated using sparse pedigrees or low-resolution molecular markers. The transition to genomic metrics of inbreeding offers unprecedented precision in quantifying realized homozygosity and its fitness costs. We conducted a systematic review and multi-level meta-analysis of 91 effect sizes from 20 wild vertebrate populations to provide a global benchmark for genomic inbreeding depression. Our results confirm a pervasive and significant negative relationship between fitness and genomic inbreeding. Males exhibited significantly stronger inbreeding depression than females. Fitness costs were consistent across life stages (developmental, adult and lifetime) and across types of fitness traits (survival vs. reproduction). We found no significant association between the magnitude of inbreeding depression and IUCN conservation status or historical isolation (discretely measured as “isolated” or “non-isolated”). As we enter a genomics era that will provide realised estimates of inbreeding, future studies can be added to this meta-analysis to provide a more comprehensive view of inbreeding depression and potentially identify patterns pertinent to evolutionary biology and conservation science.

## Introduction

Inbreeding depression, the reduction in fitness of offspring resulting from the mating of related individuals, is a key theme in evolutionary and conservation biology and a concern in conservation science. In inbred offspring, genomic segments become homozygous by descent (Charlesworth & Willis 2009) increasing the expression of deleterious recessive alleles and reducing the heterozygote advantage of overdominant loci. While inbreeding is a natural process in many social systems, anthropogenic activities have increased its frequency by forcing wild populations into demographic bottlenecks. Following bottlenecks, reduced population size is potentially compounded by inbreeding depression reducing fitness, and genetic drift reducing allelic diversity, to further accelerate population decline (Lande 1988).

Our current understanding of inbreeding depression in the wild is potentially constrained by methodological limitations. Early studies relied on pedigree analysis to calculate the expected coefficient of inbreeding (*F*), which represents the probability that two alleles are homozygous-by-descent (HBD). However, pedigrees are rarely available for wild populations, *F* can only be calculated relative to founding individuals, and it does not account for the stochastic nature of Mendelian sampling and recombination (Kardos *et al*. 2016; Wang 2016). The era of heterozygosity–fitness correlations often utilized a handful of genetic markers, such as microsatellite loci, as proxies of the genome-wide state, but these lacked statistical power regarding both genome coverage and sample size required to detect subtle fitness costs (Balloux *et al*. 2004; Slate *et al*. 2004). We now enter an era of genomics, where high-density Single Nucleotide Polymorphism (SNP) arrays and whole-genome sequencing has fundamentally shifted this landscape to enable accurate measures of inbreeding.

While the genomic study of inbreeding depression has been used in the livestock industry, its application to wild animal populations is a more recent development. Previous meta-analyses have largely focused on pedigree data (Chan 2022) or experimental laboratory populations (Vega-Trejo *et al*. 2022), which may not accurately reflect the environmental stressors or quantify the realised individual inbreeding found in nature. Genomic datasets for wild vertebrates have proliferated since 2013 (Kardos *et al*. 2016), and there is a critical need for a contemporary synthesis to determine the true magnitude of the costs of inbreeding. Crucially, the impact of inbreeding is not expected to be uniform across all contexts, but to be moderated by intrinsic biology and extrinsic environmental pressures. Identifying whether certain life-history stages, sexes, or demographic backgrounds are more susceptible to these costs is essential for predicting which populations are most vulnerable to inbreeding depression and targeting conservation interventions. This study provides a systematic review and meta-analysis of genomic inbreeding depression in freely breeding wild or feral vertebrates. By synthesizing 91 effect sizes across 20 species, we investigate if the intensity of inbreeding depression is moderated by sex, life history, fitness measure, and demographic history. Our benchmark encourages future studies of wild populations to come forth so that patterns of inbreeding depression can be disentangled with potential application in conservation.

## Methods

### Literature search

We conducted a systematic review and meta-analysis following the Preferred Reporting Items for Systematic Reviews and Meta-Analyses (PRISMA) guidelines (Moher *et al*. 2009). An initial scoping search on 1 September 2022 identified 13 primary research papers that investigated inbreeding depression in wild populations using genomic markers. Based on these texts, we constructed a Boolean search string across five concept groups: 1) inbreeding, 2) fitness, 3) wild populations, 4) genomic measures, and 5) exclusionary terms to remove irrelevant disciplines (Table S4.1). We queried Scopus, Web of Science, ProQuest, bioRxiv, and Google Scholar (first 100 hits). We searched bioRxiv and Google Scholar as these include dissertations and preprints, to reduce publication bias. To target genomic studies of inbreeding depression we filtered for texts from 2013 in our search, as the earliest study identified in our scoping search was published in 2014. The final search, conducted on 17 June 2025, yielded 969 records: 269 in Scopus, 462 in Web of Science, 19 in ProQuest, 119 in BioRxiv and 100 in Google Scholar, including all 13 of the original scoped texts. After de-duplication using Zotero, 692 unique texts remained for screening.

### Screening

We screened titles and abstracts against a standardised flowchart of five eligibility questions (Table S4.2) in a two-stage screening process using Rayyan (Ouzzani *et al*. 2016). To ensure reproducibility, a double-blind screening of the first 60 texts was conducted and, after resolving initial ambiguities, we achieved a Cohen’s Kappa of 0.857 (Cohen 1960; Landis & Koch 1977). We included studies if they investigated freely breeding wild or feral animal populations, explicitly measured inbreeding using >1,000 genomic markers, and reported a quantitative relationship between a survival or reproductive fitness trait and inbreeding. We excluded reviews unless they provided original, previously unpublished data. Following screening, we retained 39 papers for full-text eligibility assessment (Figure S4.1).

### Data extraction

We emailed the corresponding authors of five publications for further data needed to extract seven effect sizes and received responses from the authors of two papers (Clark *et al*. 2025; Crossman *et al*. 2024) within a one-month time frame, which allowed us to include a further five effect sizes in our analysis. We also added data from Lee *et al*. (2026) at this stage as they passed all the study screening criteria (Table S4.2). Therefore, we had a final dataset of ninety-two effect sizes from twenty-seven papers to be included in the meta-analysis (Table S4.3). To maintain statistical independence when studies reported multiple inbreeding estimators for the same population and trait, we applied a predefined hierarchy (Table 4.1).

**Table 4.1.** Hierarchy of genomic inbreeding estimates where studies reported multiple inbreeding depression estimates.

| Rank | Genomic inbreeding coefficient | Explanation |
| --- | --- | --- |
| 1 | Runs of homozygosity ($F_{ROH}$) | Quantifies continuous homozygous segments as a close proxy to HBD. Where multiple length thresholds were reported, the longest $F_{ROH}$ was prioritized to capture recent parental relatedness. |
| 2 | Genomic relatedness matrix ($F_{GRM}$) | An allele-frequency-weighted measure of genome-wide homozygosity. It prioritizes homozygosity for rare alleles, which are more likely to be HBD and deleterious (VanRaden 2008). |
| 3 | Correlation of uniting gametes ($F_{UNI}$) | Estimates the correlation between uniting parental gametes, accounting for the variance in allele frequencies across the genome (Yang *et al.* 2011). |
| 4 | Excess homozygosity ($F_{HOM}$) | Calculates the observed number of homozygous loci relative to the expectation under Hardy-Weinberg Equilibrium. This represents a standardized count of homozygotes but, unlike $F_{GRM}$, does not weight by allele rarity (Purcell *et al.* 2007). |
| 5 | $sMLH$ | The total number of heterozygous loci divided by the population mean, standardized by the number of typed markers. This correction accounts for differential marker coverage across individuals (Stoffel *et al.* 2016). |
| 6 | $H_o$ | The unstandardized genome-wide proportion of homozygous loci. This is a measure of identity-by-state (IBS) and serves as a fundamental proxy for genetic diversity (Chapman *et al.* 2009). |

We prioritized *F*_ROH_ as the most direct measure of HBD, followed by allele-frequency-weighted matrices (*F*_GRM_, then *F*_UNI_, then *F*_HOM_) and, finally, absolute measures of individual homozygosity (*sMLH* then *H*_O_). All primary results were converted into a standardized effect size: Pearson’s correlation coefficient (*r*). A negative *r* was defined as a signal of inbreeding depression (decreased fitness associated with increased inbreeding).

We recorded the following data: effect size ID, study ID, population ID, author, year, publication type, title, species name (common), species binomial name, taxon, conservation status (IUCN definitions), sample size, genomic inbreeding coefficient used in analysis, inbreeding coefficient grouped (runs of homozygosity (*F*_ROH_), excess homozygosity (*F*_GRM_*, F*_UNI_, or *F*_HOM_) or individual homozygosity (*sMLH* or *H*_O_)), number of markers used, sex measured (males, females, or grouped), fitness metric type (survival or reproductive), specific fitness metric, lifestage (for survival traits, whether they could be classed as developmental, lifetime or adult), effect size type for conversion to correlation coefficient *r*, raw values used to calculate effect size, where in the text effect sizes can be found, direction of *r*, correlation coefficient *r*, Fisher’s *z* and Variance. We used the results or data presented after any exclusion of outliers.

We implemented a systematic workflow to convert all primary study outcomes into a standardized effect size, Pearson’s correlation coefficient (*r*), representing the strength and direction of the relationship between fitness and genomic inbreeding across both continuous and categorical data. We defined a negative *r* as a signal of inbreeding depression, where increased inbreeding was associated with decreased fitness.

Effect size conversions to *r* (Altman & Bland 2011; Borenstein *et al*. 2009; Nakagawa & Cuthill 2007; Nakagawa & Schielzeth 2013; Rosenberg *et al*. 2013):

1. Pearson’s *r* : Where reported, Pearson’s *r* and the associated sample size (*n*) were extracted directly.
2. Cohen’s *d* : For studies comparing discrete groups, we calculated Cohen’s *d* from group means and standard deviations (or standard errors) as follows :

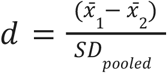

and converted it to *r* using the formula:

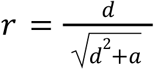

where *a* is a correction factor for unequal group sizes:

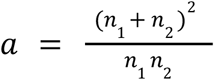
3. Test statistics (*t*, *F*, and *x^2^*):
*t*-statistics were converted to *r* using:

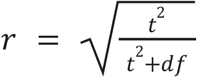

*F*-statistics (where *df* = 1) were converted using:

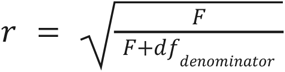

Chi-square values (*df* = 1) were converted via:

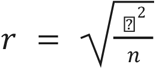
4. Regression coefficients (*β*): For studies reporting linear models, we employed a *t*-statistic pathway. Unstandardized coefficients were converted to *t* using:

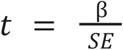

If the standard error (*SE*) was not reported, it was derived from 95% confidence intervals as:

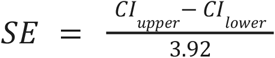

or back-calculated from exact *p*-values by first determining the corresponding *z*-score using the inverse standard normal cumulative distribution (Altman & Bland 2011):

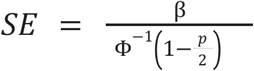 Standardized *β* coefficients were treated as approximations of *r* if both the predictor and response variables were standardized.
5. Log-odds ratios (*L*): Log-odds ratios from generalized linear models were converted to Cohen’s *d* via the transformation before converting to *r*:

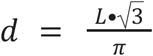

We transformed all *r* values to Fisher’s *z* to normalize the sampling distribution:

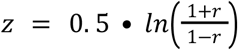

The associated sampling variance (*V_z_*) for each effect size was calculated as:

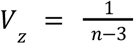

If *n* was not reported, we estimated it as *df* + *k* + 1, where *k* was the number of fixed predictors.

### Analysis

We performed analyses in R v.4.4.3 using the rma.mv function in the metafor package (Viechtbauer 2010). We removed one effect size (*z* = −1.73) as an outlier, which was likely to be a data artifact as the standard deviation was inconsistently larger than for similar models in the study by an order of magnitude. This resulted in a final dataset of *k* = 91 effect sizes.

We first fitted an intercept-only model to estimate the global magnitude of genomic inbreeding depression. We assessed total heterogeneity using Cochran’s *Q* test and calculated the total *I*^2^ statistic to quantify the proportion of observed variance due to true heterogeneity, rather than sampling error.

To explore sources of variation in inbreeding depression, we conducted multi-level meta-regressions (mixed-effects models), including specific fixed-effect moderators. We tested the influence of:

1. Sex (male, grouped or female)
2. Lifestage (developmental, adult or lifetime, with developmental traits being those measured in the pre-reproductive stage for that species, adult traits those measured during the reproductive stage for that species, and lifetime traits those measured at repeated intervals throughout the lifetime)
3. Fitness trait (survival or reproduction)
4. IUCN Conservation Status (Least Concern, Vulnerable, Endangered, Critically Endangered or Data Deficient, with feral Sable Island horses and Soay sheep categorised as Least Concern)
5. Historical isolation (isolated or non-isolated, with isolated populations defined as island or restricted, regional endemics and non-isolated populations as being historically broadly distributed across mainland continents or oceans)
6. Genomic inbreeding coefficient used (runs of homozygosity (*F*_ROH_), excess homozygosity (*F*_GRM_*, F*_UNI_, or *F*_HOM_) or individual homozygosity (*sMLH* or *H*_O_))

To control for non-independence, we included ‘Study ID’ (accounting for multiple estimates within the same publication), ‘Population ID’ (accounting for multiple estimates derived from the same biological population), and ‘Phylogeny’ (the phylogenetic variance–covariance matrix, accounting for phylogenetic non-independence) as random effects in all models. We created the phylogenetic variance–covariance matrix from all species in the dataset using the Open Tree of Life taxonomy (Michonneau *et al*. 2016) and median divergence times (Figure S4.2), and assuming a Brownian motion model of evolution in the ape package (Paradis & Schliep 2019). We noted a high degree of collinearity between Study ID and Phylogeny, with Study ID subsuming nearly all phylogenetic signal (*σ^2^*_study_ = 0.015; *σ^2^*_phylo_ < 0.001) because most studies concerned a single taxon, but we chose to retain Study ID to control for pseudoreplication. For categorical moderators, we assessed significance using omnibus Wald-type tests (*Q*_M_). To visualise variation across taxa (Figure S4.3), we fitted a separate model with Species as a fixed factor without an intercept to estimate the mean inbreeding depression (*z*) and associated 95% confidence intervals for each species independently. We visualised results using the orchaRd package (Nakagawa *et al*. 2021). We assessed the potential for publication bias visually using contour-enhanced funnel plots and statistically using Egger’s regression.

## Results

### Overall results

Across all 91 effect sizes from 27 studies (Table S4.3) in 20 species, we found a significant negative relationship between genomic fitness and inbreeding (Overall Fisher’s *z* = −0.114, 95% CI [−0.161, −0.066], p < 0.001, Figures 4.1 and S2). Effects were highly heterogeneous (*I*^2^ = 85.3%). All species were vertebrates and mostly comprised mammals (*n* = 10) and birds (*n* = 8), but also non-avian reptiles (*n* = 1) and fish (*n* = 1).

**Figure 4.1.**
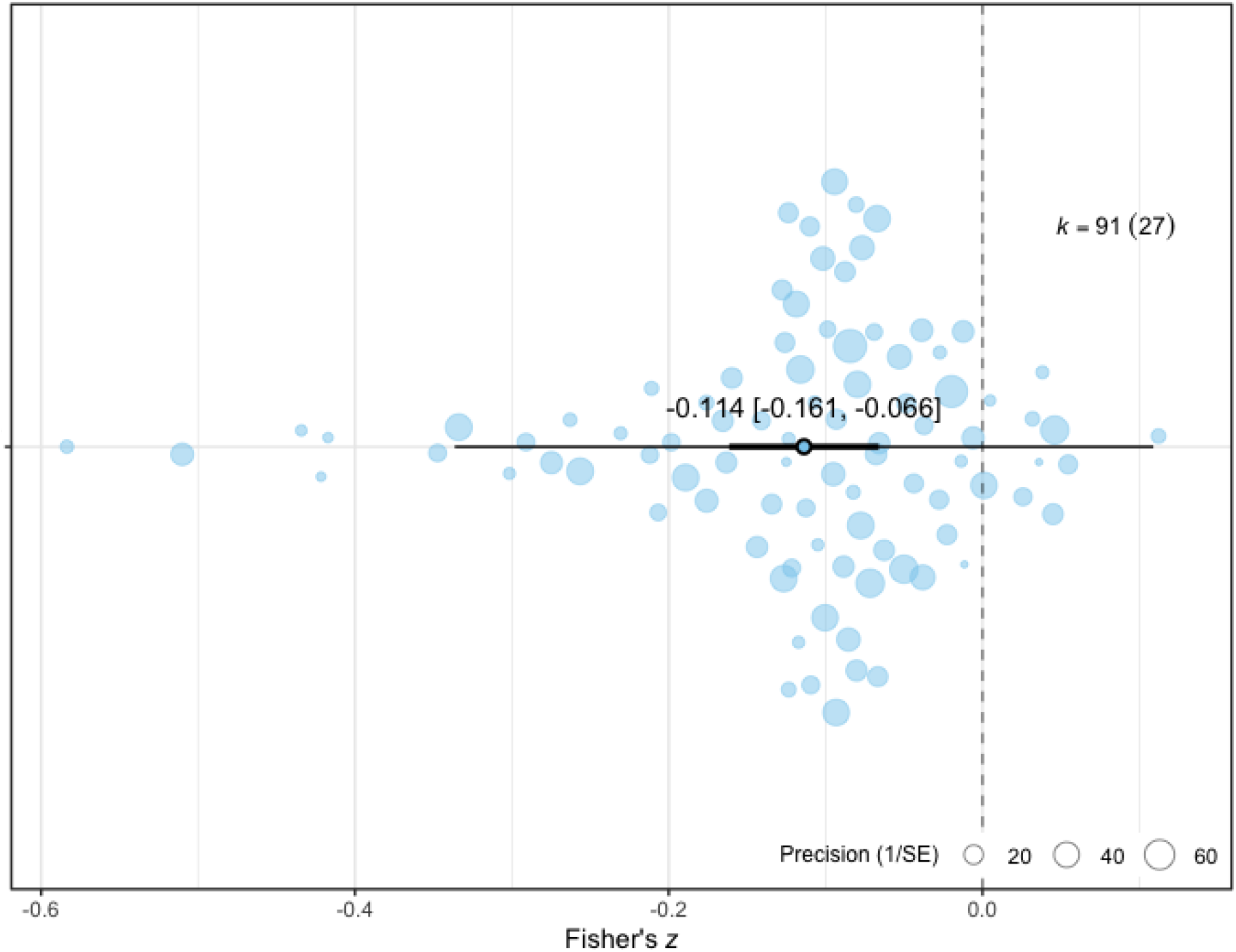
The overall magnitude of genomic inbreeding depression using 91 effect sizes (*k*) from 27 studies in 20 wild animal species. The plot displays the marginal mean estimate (large point) with the 95% confidence interval (thick bar) and 95% prediction interval (thin whiskers) derived from a multi-level meta-regression accounting for study identity, population identity, and phylogeny as random effects. Individual data points are jittered and scaled by precision.

### Sex effect

Sex significantly affected inbreeding depression (*Q*_M_ = 8.06, df = 2, *p* = 0.018). Males exhibited significantly higher levels of inbreeding depression compared to females (*β*_diff_ = 0.039, 95% CI [−0.041, −0.037], *p* = 0.011, Figure 4.2).

**Figure 4.2.**
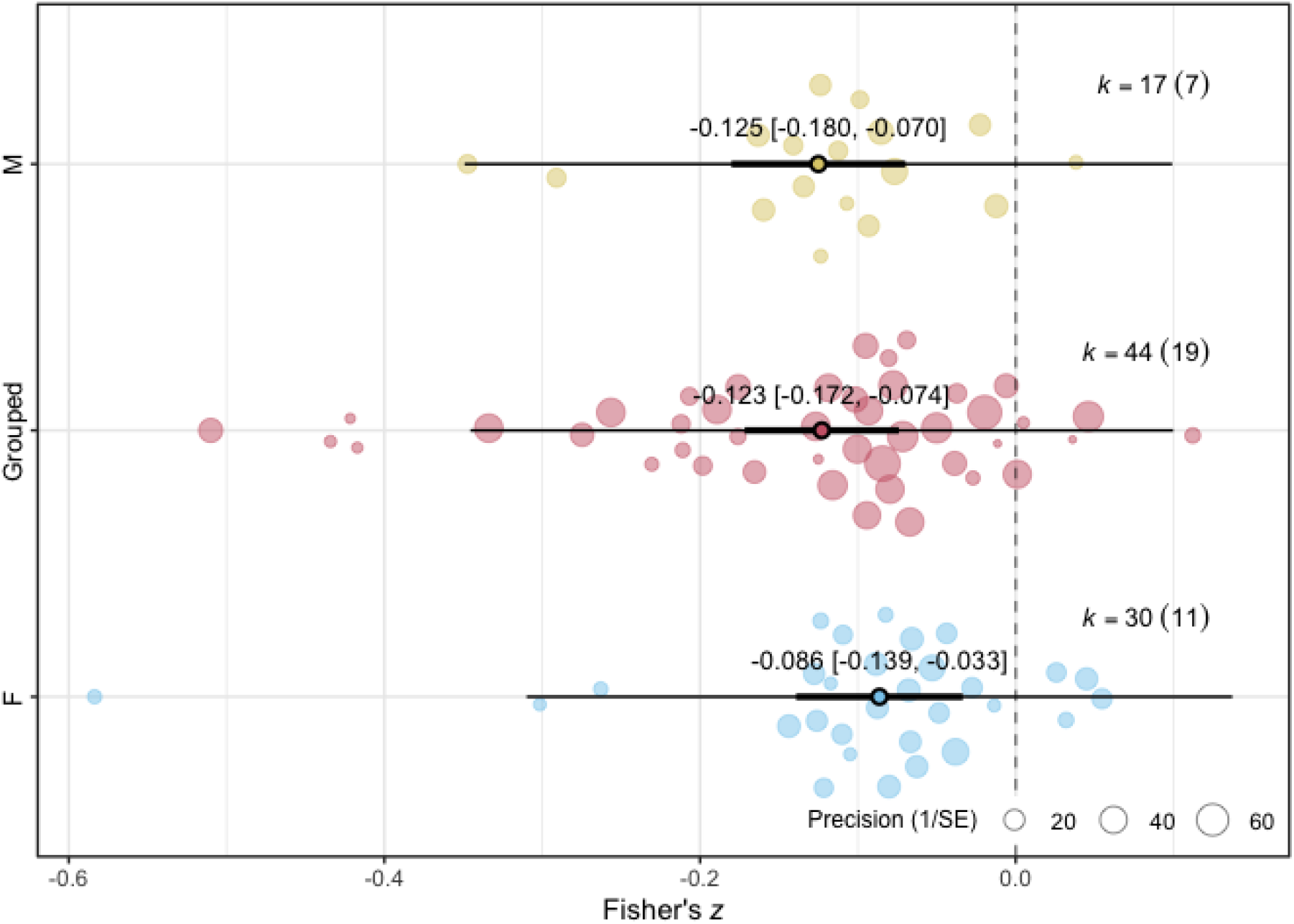
Sex effect on genomic inbreeding depression using 91 effect sizes (*k*) from 27 studies and 20 wild animal species. The plot displays the marginal mean estimates (large points) with the 95% confidence intervals (thick bars) and 95% prediction intervals (thin whiskers) for males (M), females (F) and pooled sex data (Grouped). derived from a multi-level meta-regression accounting for study identity, population identity and phylogeny as random effects. Individual data points are jittered and scaled by precision.

### Lifestage effect

Inbreeding depression was similar across all lifestages (*Q*_M_ = 0.186, df = 2, *p* = 0.911, Figure 4.3).

**Figure 4.3.**
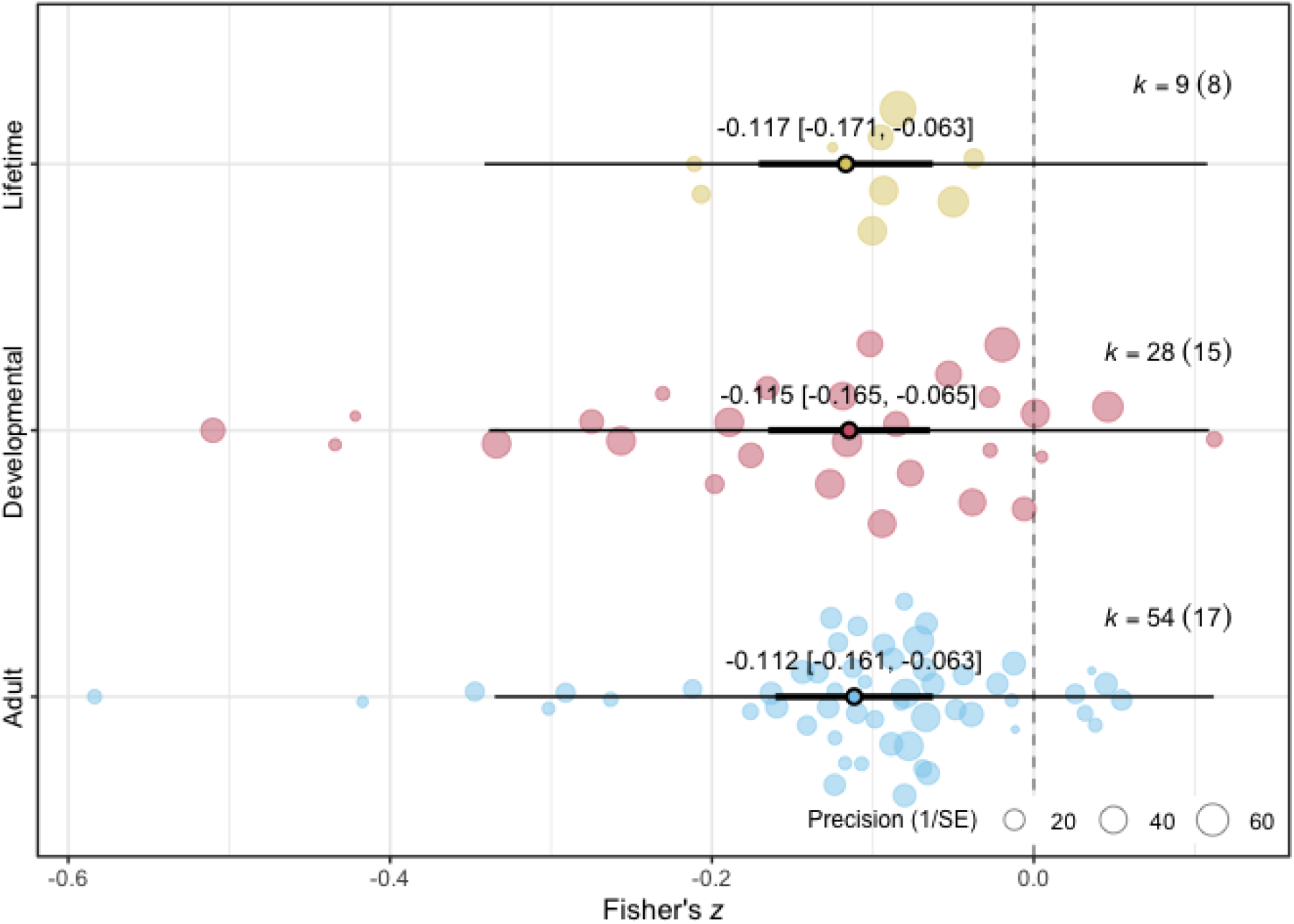
Lifestage effect on genomic inbreeding depression using 91 effect sizes (*k*) from 27 studies and 20 wild animal species. The plot displays the marginal mean estimates (large points) with the 95% confidence intervals (thick bars) and 95% prediction intervals (thin whiskers) for lifetime, developmental and adult fitness traits, derived from a multi-level meta-regression accounting for study identity, population identity, and phylogeny as random effects. Individual data points are jittered and scaled by precision.

### Fitness effect

The type of fitness trait, measured as “survival” or “reproduction”, did not significantly moderate the magnitude of inbreeding depression (*Q*_M_ = 0.082, df = 1, *p* = 0.775, Figure 4.4).

**Figure 4.4.**
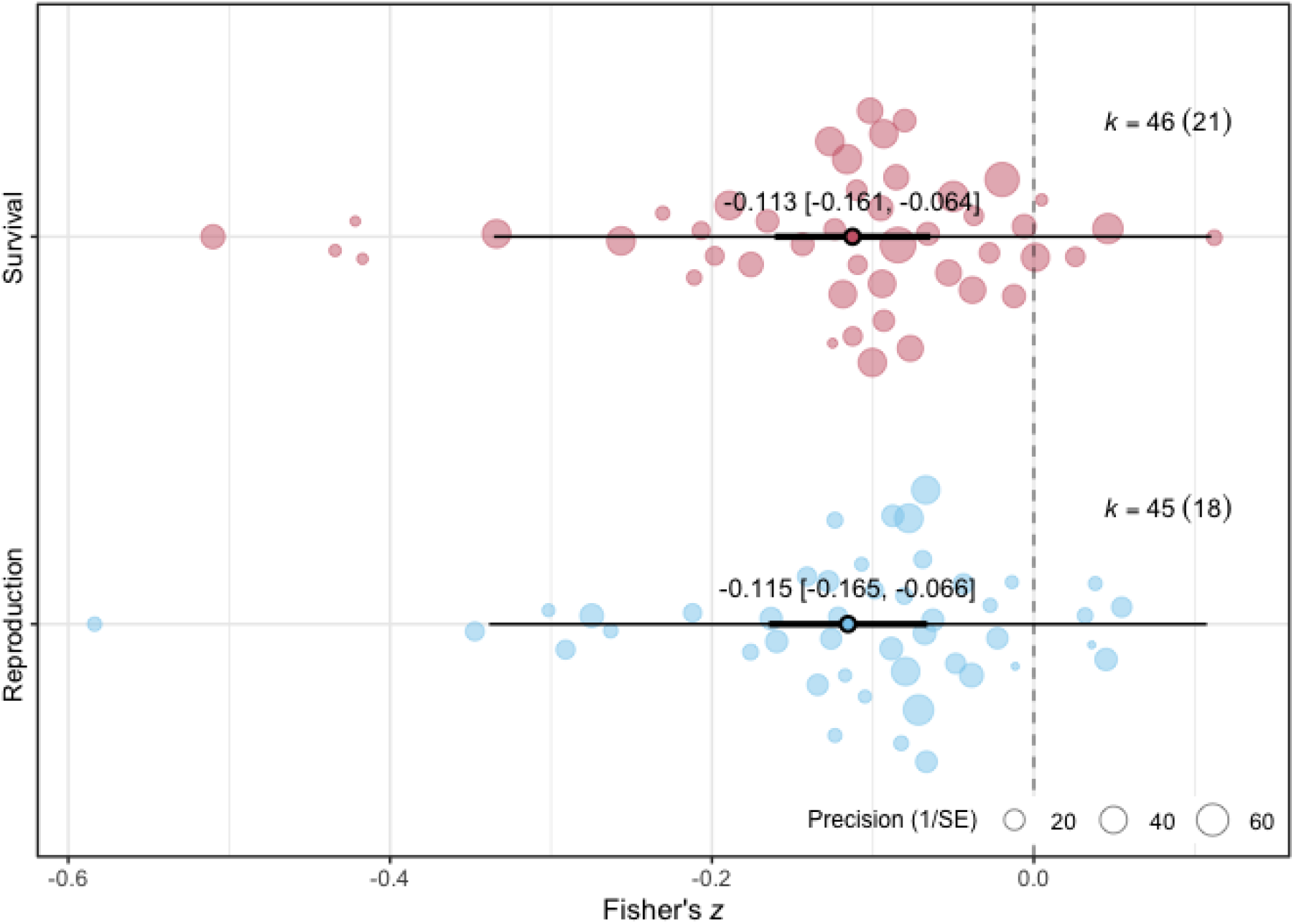
Effect of fitness trait measured on genomic inbreeding depression using 91 effect sizes (*k*) from 27 studies and 20 wild animal species. The plot displays the marginal mean estimates (large points) with the 95% confidence intervals (thick bars) and 95% prediction intervals (thin whiskers) for survival and reproduction fitness traits derived from a multi-level meta-regression accounting for study identity, population identity, and phylogeny as random effects. Individual data points are jittered and scaled by precision.

### Conservation status effect

Species conservation status, as categorised by the IUCN to be of least concern, vulnerable, endangered, critically endangered or data deficient, did not significantly moderate the magnitude of inbreeding depression (*Q*_M_ = 2.001, df = 4, p = 0.734, Figure 4.5).

**Figure 4.5.**
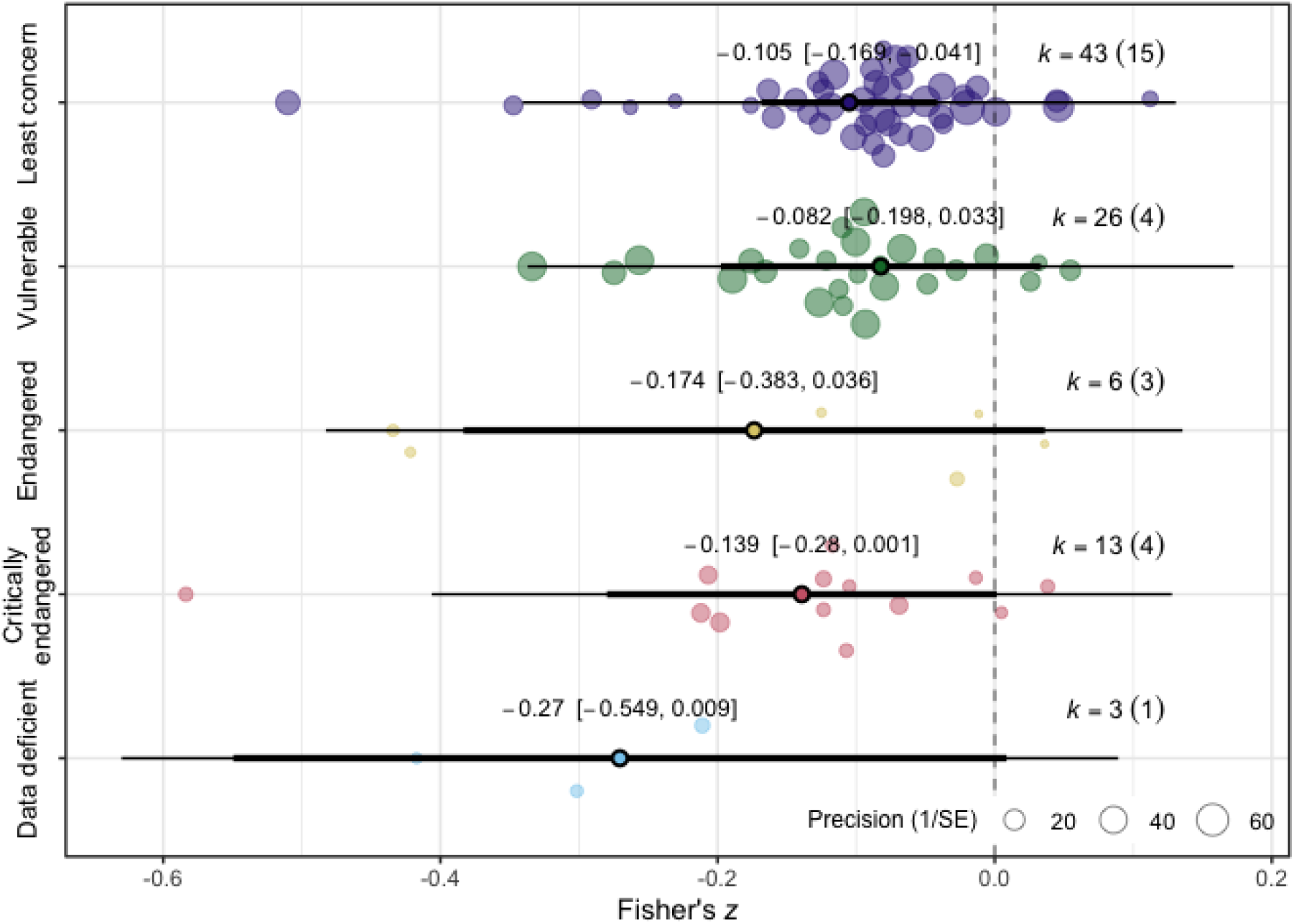
Effect of species IUCN conservation status on genomic inbreeding depression using 91 effect sizes (*k*) from 27 studies and 20 wild animal species. The plot displays the posterior mean estimates (large points) with the 95% confidence intervals (thick bars) and 95% prediction intervals (thin whiskers) for each IUCN category derived from a multi-level meta-regression accounting for study identity, population identity, and phylogeny as random effects. Individual data points are jittered and scaled by precision.

### Historical isolation effect

Historical isolation, categorised as “isolated” or “non-isolated”, did not significantly moderate the magnitude of inbreeding depression (*Q*_M_ = 0.495, df = 1, *p* = 0.482, Figure 4.6).

**Figure 4.6.**
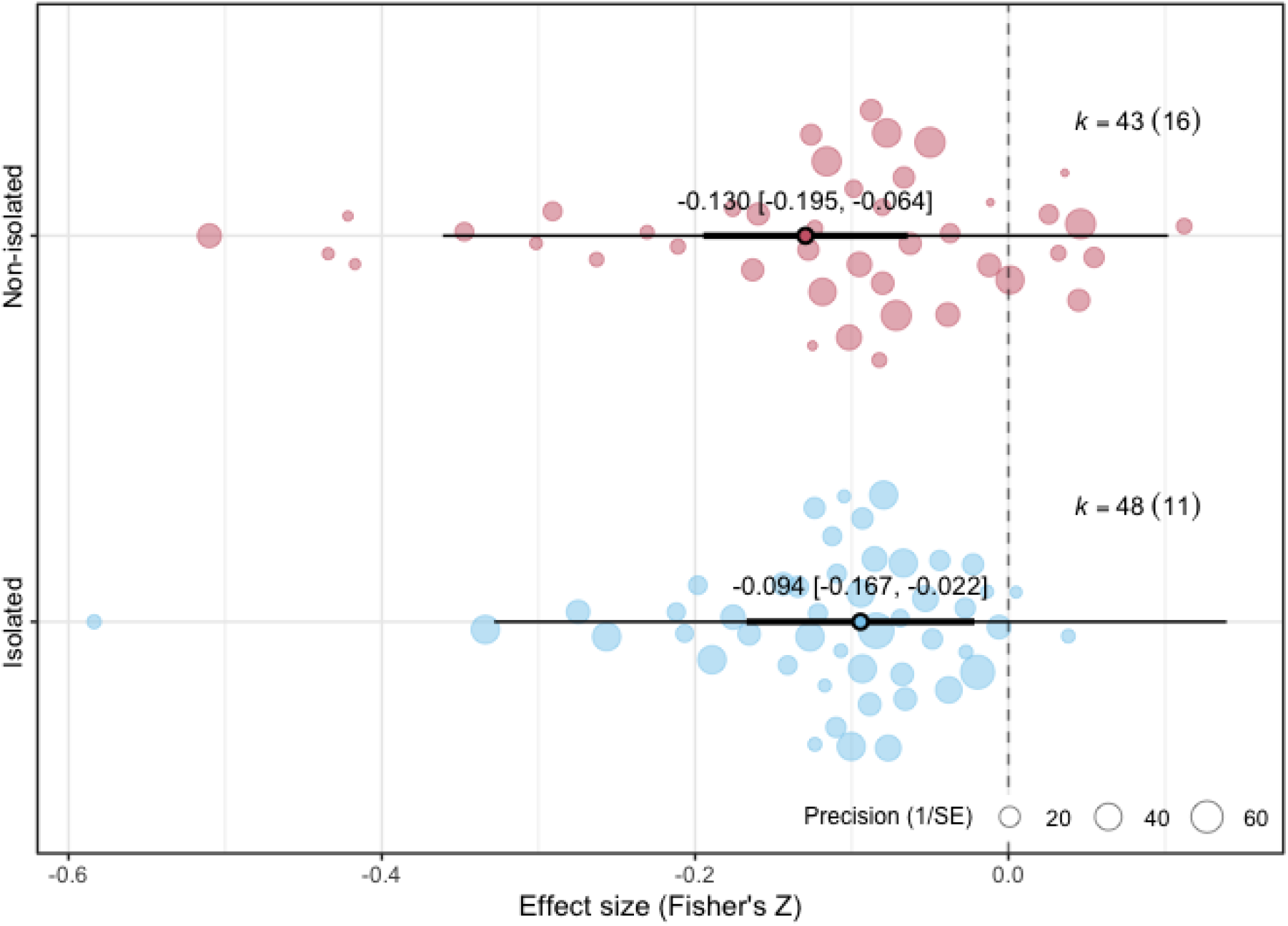
Effect of historical isolation on genomic inbreeding depression using 91 effect sizes (*k*) from 27 studies and 20 wild animal species. The plot displays the posterior mean estimates (large points) with the 95% confidence intervals (thick bars) and 95% prediction intervals (thin whiskers) relative to population isolation, derived from a multi-level meta-regression accounting for study identity, population identity, and phylogeny as random effects. Individual data points are jittered and scaled by precision.

### The effect of the genomic inbreeding coefficient

The method of estimating genomic inbreeding did not significantly moderate the magnitude of inbreeding depression (*Q*_M_ = 0.060, df = 2, *p* = 0.970, Figure 4.7).

**Figure 4.7.**
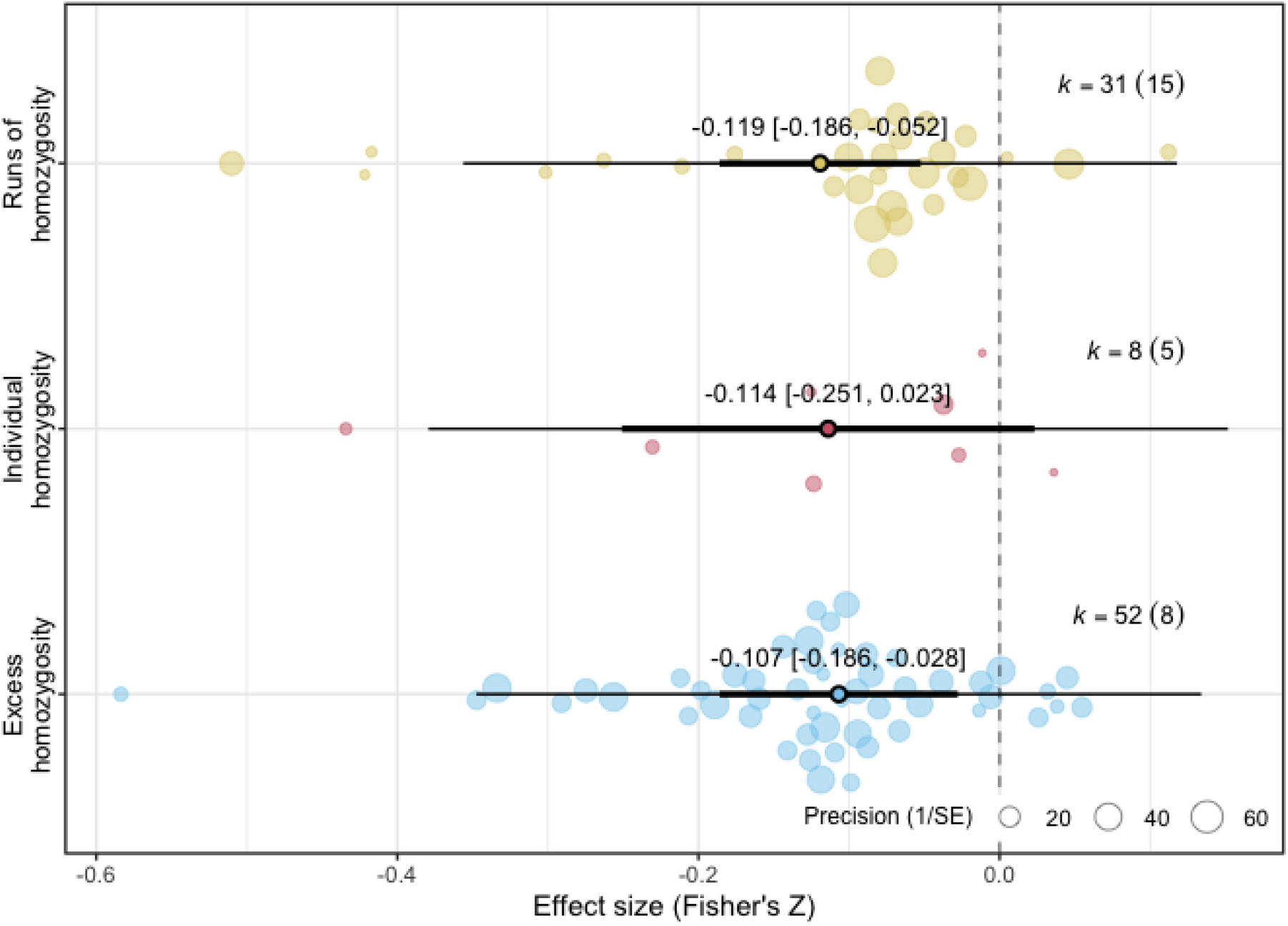
Effect of genomic inbreeding coefficient used on genomic inbreeding depression using 91 effect sizes (*k*) from 27 studies and 20 wild animal species. The plot displays the posterior mean estimates (large points) with the 95% confidence intervals (thick bars) and 95% prediction intervals (thin whiskers) for alternative coefficients of inbreeding derived from a multi-level meta-regression accounting for study identity, population identity, and phylogeny as random effects. Individual data points are jittered and scaled by precision.

### Publication bias

Visual inspection of the funnel plot revealed a generally symmetrical distribution of effect sizes (Figure S4). Consistent with this, the modified Egger’s test indicated no significant relationship between the effect size and its sampling error (*Q*_M_ = 0.031, df = 1, *p* = 0.860), suggesting that our results are not driven by the preferential publication of significant results from smaller studies.

## Discussion

Our meta-analysis confirms a pervasive negative relationship between fitness and genomic inbreeding across 20 wild vertebrate populations (*z* = −0.114). Our measured fitness cost in wild vertebrates is greater than found in a previous meta-analysis of 43 effect sizes (Chan 2022), which measured the fitness difference between pedigree-based comparisons of inbred and outbred groups. This suggests that inbreeding costs may have been previously underestimated using pedigree-based metrics by not capturing the realised variance in homozygosity created by Mendelian sampling (Kardos *et al*. 2016; Wang 2016). Conversely, our genomic estimate is more conservative than in a recent meta-analysis of inbreeding studies in animals (β [95% CI] = 0.38 [0.06; 0.7], or *z* = 0.189), which compared highly inbred groups with outbred controls studied under experimental conditions (Vega-Trejo *et al*. 2022). While such experiments maximize statistical power to detect a cost, they may not reflect the continuous gradients of inbreeding found in nature. As sequencing becomes faster and more affordable, access to continuous, genomic measures of inbreeding may provide more precision, to capture the realised costs of inbreeding depression in wild populations.

As genomic studies of inbreeding depression use the autosomal quantification of inbreeding, our finding that inbreeding depression is significantly less strong in females might be explained by sexual selection (Grieshop *et al*. 2021; Janicke *et al*. 2013; Mallet & Chippindale 2011). In many animal populations, male fitness exhibits greater variance due to the expression of costly sexually selected traits. Sexually selected male traits are potentially more sensitive to the effects of deleterious, recessive alleles being exposed in the homozygous state, leading to a higher detectable cost of inbreeding in males (Enders & Nunney 2010). As genomic estimates of inbreeding are mostly obtained from autosomal data and half the species tested were mammals (XY males are heterogametic), whereas nearly half were birds (ZZ males are homogametic), the sex effect is not likely explained by heterogametic exposure of recessive alleles (Ebel & Phillips 2016). Future work could re-visit this analysis by investigating how inbreeding depression is moderated by a proxy measure of sexual selection that is comparable across species, such as the difference in reproductive variance between the sexes.

Our results show that the fitness costs of inbreeding are consistent across lifestages (developmental, adult or lifetime) and trait types (survival vs. reproduction). This suggests that the genetic load consists of a broad spectrum of deleterious alleles that manifest throughout the entire life cycle. Lifestage analysis is subject to survivorship bias as the most severe deleterious effects are likely purged through embryonic or neonatal mortality before individuals can be sampled in the wild. We also acknowledge our categorisations of fitness by lifestage and trait type may be too broad. For example, grouping first-year, annual, and adult survival under a single “survival” trait, or fledgling and first-year survival under a “developmental” lifestage, may overlook stage-specific sensitivities to inbreeding that vary significantly across vertebrate life histories. Continued long-term monitoring of wild populations will help understand the impact of inbreeding across lifestage, as currently, such patterns appear species-specific (Auclair *et al*. 2025; Beccardi *et al*. 2024; Charlesworth & Hughes 1996; Huisman *et al*. 2016; Keller & Waller 2002; Trask *et al*. 2021).

We found no significant differences in the magnitude of inbreeding depression across IUCN conservation status categories. This may be because IUCN criteria prioritize contemporary demographic trends and geographic ranges and do not specifically account for the quantification of masked genetic load. In ‘Critically endangered’ island-endemic kākāpōs (*Strigops habroptilus*) of New Zealand, a long history of small population size provided the opportunity for natural selection to purge the genetic load (Dussex *et al*. 2021), resulting in a relatively small effect size for genomic inbreeding depression in this species (Foster *et al*. 2021). Conversely, the, ‘Least concern’ red deer population (*Cervus elaphus*) on the Isle of Rum, which descend from multiple reintroductions and translocations from diverse UK and European sources (Nussey *et al*. 2006) exhibit relatively strong inbreeding depression ((Hasik *et al*. 2025; Hewett *et al*. 2025), Figure S4.3). These source populations, if historically large, would carry masked genetic load, which had not previously been exposed until high inbreeding environments within the Isle of Rum. This conclusion may be refuted by our finding that historically “isolated” and “non-isolated” populations experience similar inbreeding depression. However, our quantification of potential for masked genetic load is limited by using two discrete categories of historical isolation that are potentially subjective. Future meta-studies could study moderation of inbreeding depression by metrics of genetic load that are easily comparable and at a continuous scale. (Kyriazis *et al*. 2025) describe one such metric of inbreeding depression potential, defined as *F_ROH_*, relative to mean heterozygosity in non-ROH regions.

We found little effect of the type of genomic inbreeding coefficient measured on the magnitude of inbreeding depression, provided marker density is sufficiently high (> 1000s SNPs). Although the different genomic estimators used across our sampled studies are not strictly equivalent in their absolute calculation, they are suitable for meta-analysis because the calculation of a correlation coefficient (*r* or *z*) depends primarily on the relative ranking of individuals by their degree of inbreeding. Previous empirical comparisons have demonstrated that when marker density is high, different genomic inbreeding metrics are strongly positively correlated (Alemu *et al*. 2020; Forutan *et al*. 2018). As our analysis focuses on the strength of the relationship between inbreeding and fitness, the rank-order consistency between these metrics ensures that our global effect size remains a robust and reliable estimate of inbreeding depression. However, *F_ROH_* is often considered the most precise measure of realized inbreeding by directly quantifying segments of the genome that are HBD, especially in small, unstructured populations that are often sampled in wild species (Caballero *et al*. 2021, 2022; Kardos *et al*. 2016; Lavanchy *et al*. 2024; Nietlisbach *et al*. 2019; Peripolli *et al*. 2017). From the studies sampled in our review, only those that use *F_ROH_* enable identification of the specific evolutionary mechanisms driving those correlations, through parameterisation of varying ROH lengths that distinguish ancient from recent inbreeding and by pinpointing regions of the genome that contribute to inbreeding depression (Duntsch *et al*. 2023; Hasselgren *et al*. 2018; Stoffel *et al*. 2021). Sometimes calling ROH can be constrained by requiring high-density SNP data (Duntsch *et al*. 2021; Lavanchy & Goudet 2023) and by not capturing rare deleterious mutations that are purged under demographic histories of large effective population size (Lavanchy *et al*. 2024; Yengo *et al*. 2017). Issues of SNP-density can be mitigated through high-density genotype imputation of low-coverage sequencing data, a strategy successfully employed in livestock (Ablondi *et al*. 2023; Martikainen *et al*. 2018; Mota *et al*. 2024) and more recently in wild species (Lee *et al*. 2026; Stoffel *et al*. 2021). While the sensitivity of genomic estimators to population structure remains a concern, the implementation of mixed models incorporating a pedigree or genomic relatedness matrix provides a robust solution. By accounting for the non-independence of observations, these models effectively isolate inbreeding effects from background genetic structure, ensuring that fitness correlations, even those driven by rare deleterious alleles, can be reliably estimated (Lavanchy *et al*. 2024).

Despite the global scope of this synthesis, our analysis is constrained by the relatively small number of populations currently available in the genomic literature (*k* = 91 from 20 populations). Limited sample size, combined with the substantial unexplained heterogeneity observed in our models (*^2^_tota_*_l_= 85.3%), suggests that the realised global mean of inbreeding depression is still subject to refinement. The long-term studied populations not identified by our systematic review are primed for application of genomic data (Bonnet *et al*. 2022; Culina *et al*. 2021) to refine our global understanding of inbreeding depression in the wild.

Furthermore, our dataset is heavily biased toward mammals and birds, which may limit the generalizability of these findings to other animals. To address these limitations and facilitate future syntheses, we provide the complete R-script and PRISMA-compliant workflow used in this study as a supplementary resource. This will allow researchers to repeat this meta-analysis as more genomic inbreeding depression research continues following this systematic review (Tsujimoto *et al*. 2025).

Our results have implications for how we assess extinction risk in the genomic era. The observed male-biased inbreeding depression suggests that for species with intense sexual selection, standard demographic models may underestimate the probability of population collapse. If males are disproportionately sensitive to inbreeding, the effective population size (*N_e_*) may be lower than census counts suggest, warranting a revisit of threat categories.

Furthermore, the lack of correlation between IUCN status and inbreeding depression indicates that “Least Concern” species could harbour masked genetic load, especially following recent bottlenecks. We suggest that future meta-research following our framework could serve as a predictive tool, identifying the physiological and life-history traits that make certain species more vulnerable to inbreeding depression, allowing for a more proactive approach to global biodiversity conservation.

## Author contributions

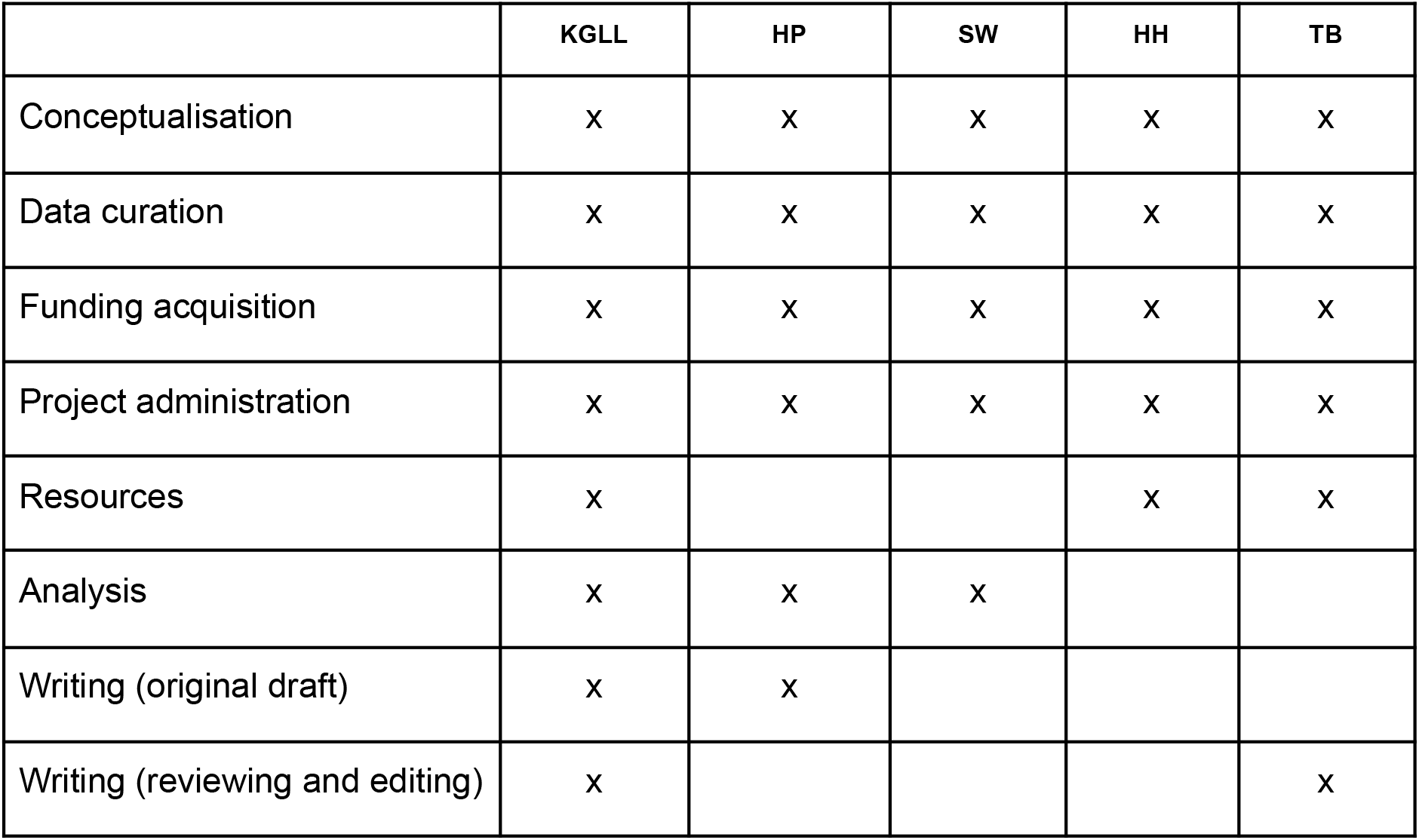

## Acknowledgements

We thank the following:

NERC grant NE/B504106/1 to T.B.

A University of Sheffield Undergraduate Research Experience (SURE) grant awarded to Hannah Parkes.

## Data availability

Scripts and data can be found at: https://github.com/kiran-lee/InbreedingDepressionWildMetaAnalysis

## Supplementary materials

**Figure S4.1.**
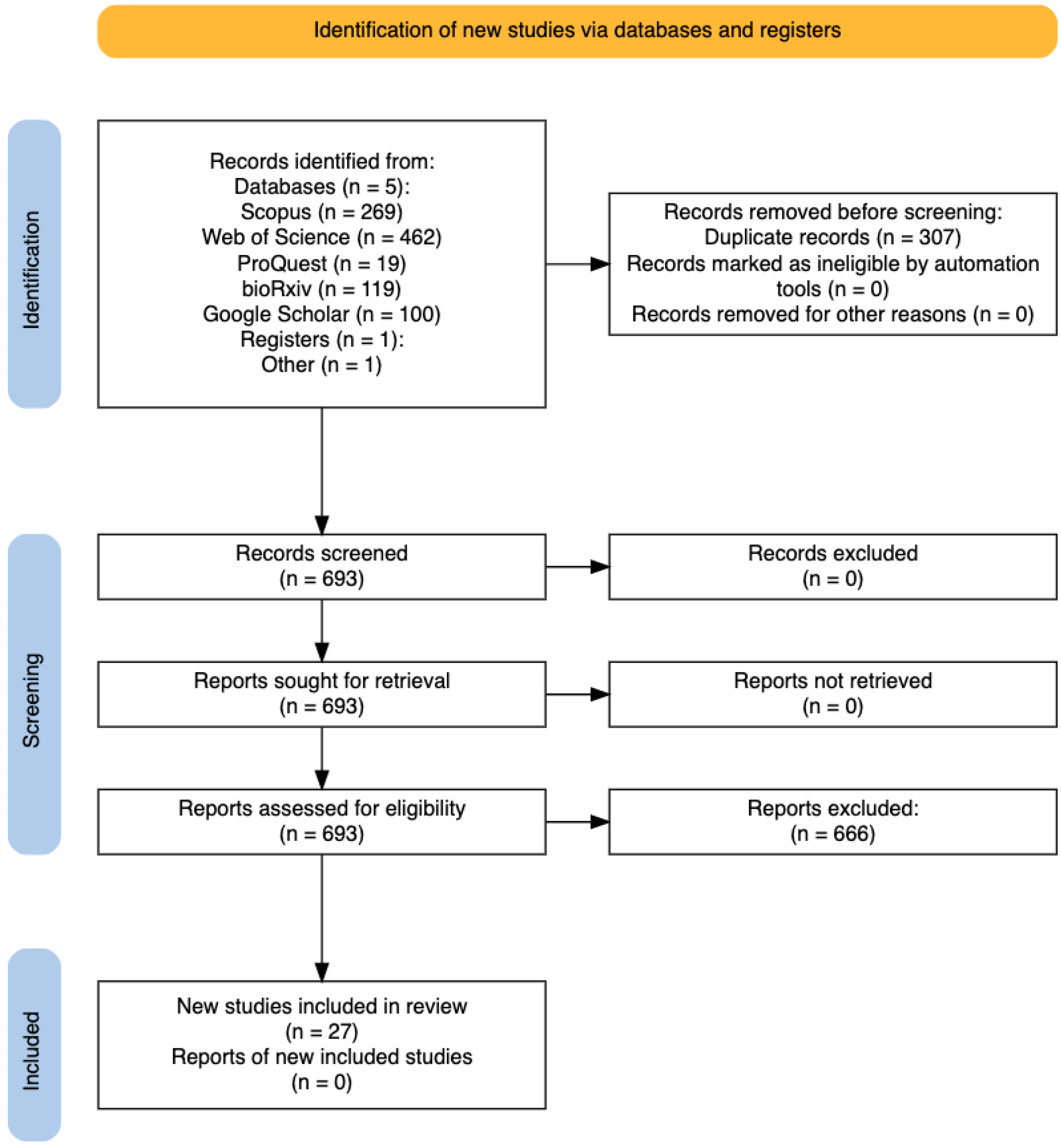
PRISMA flow diagram detailing study identification, and screening (Haddaway et al. 2022).

**Figure S4.2.**
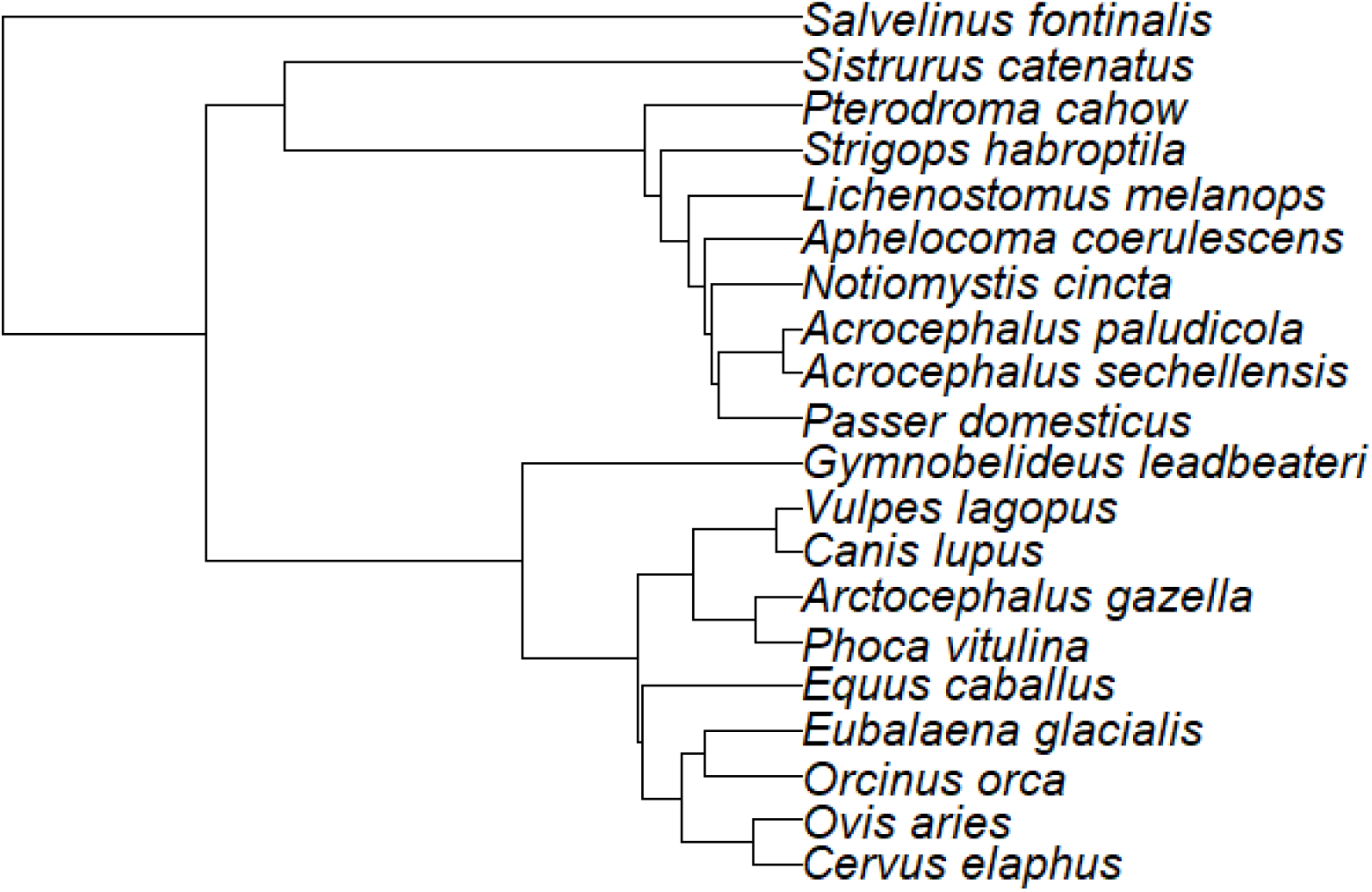
Phylogenetic tree of all 20 species, used for controlling phylogenetic independence.

**Figure S4.3.**
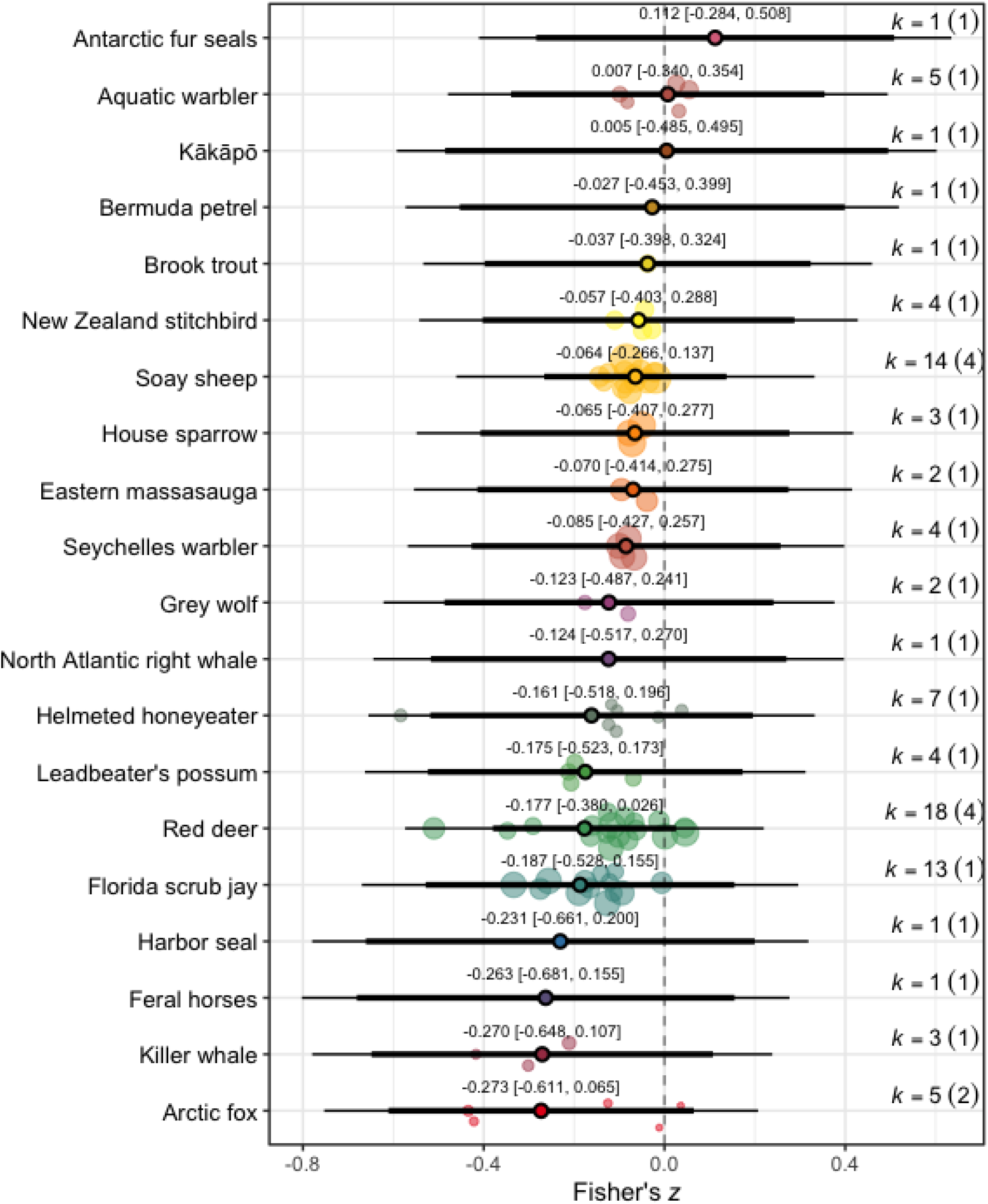
Genomic inbreeding depression estimates from 20 wild animal species using 91 effect sizes (*k*) from 27 studies. The plot displays the marginal mean estimate (large points) with 95% confidence intervals (thick bars) and 95% prediction intervals (thin whiskers) for each population derived from a multi-level meta-regression accounting for study identity, population identity, and phylogeny as random effects. Individual data points are jittered and scaled by precision.

**Figure S4.**
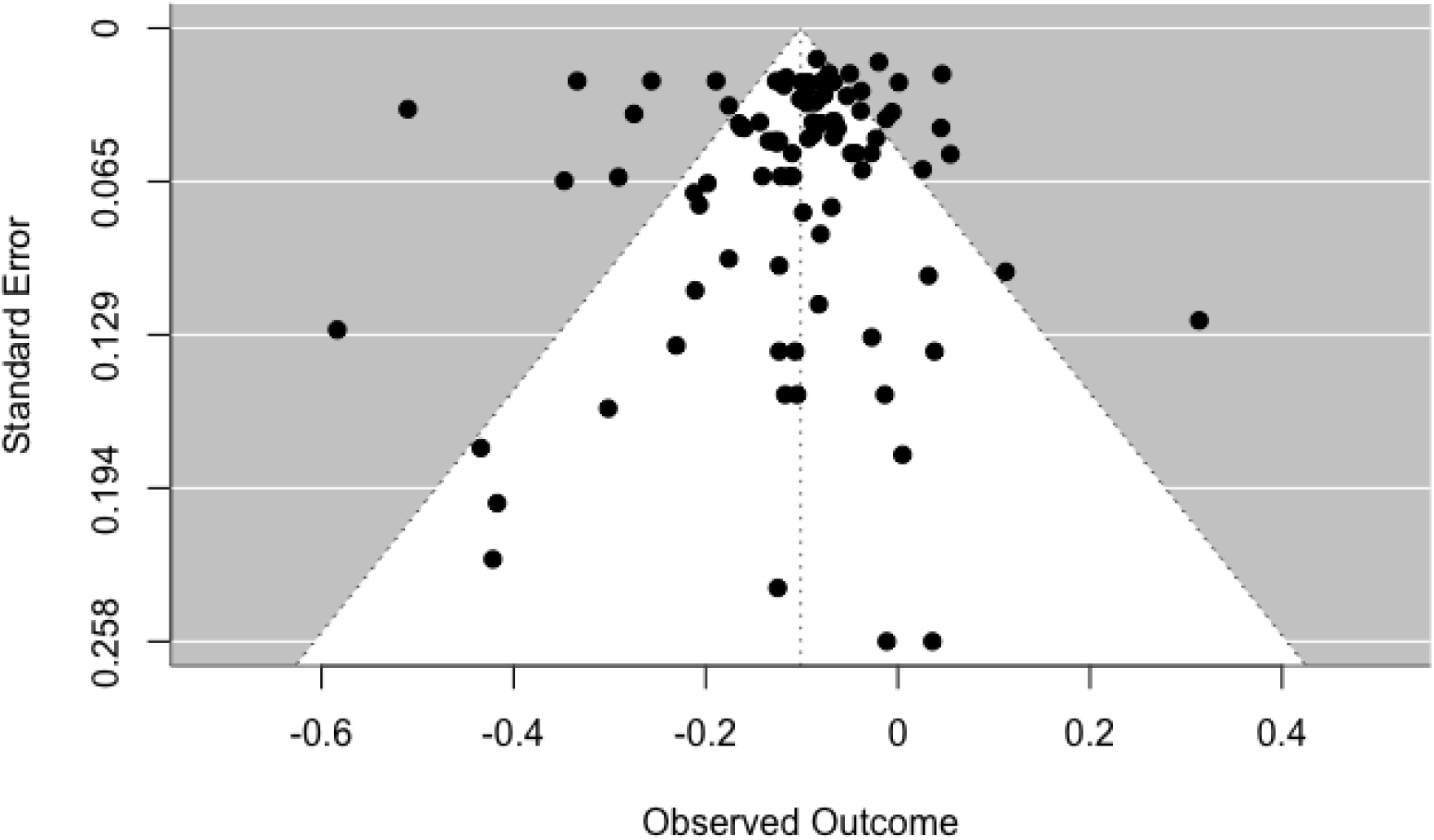
Funnel plot of all 91 effect sizes.

**Table S4.1.**
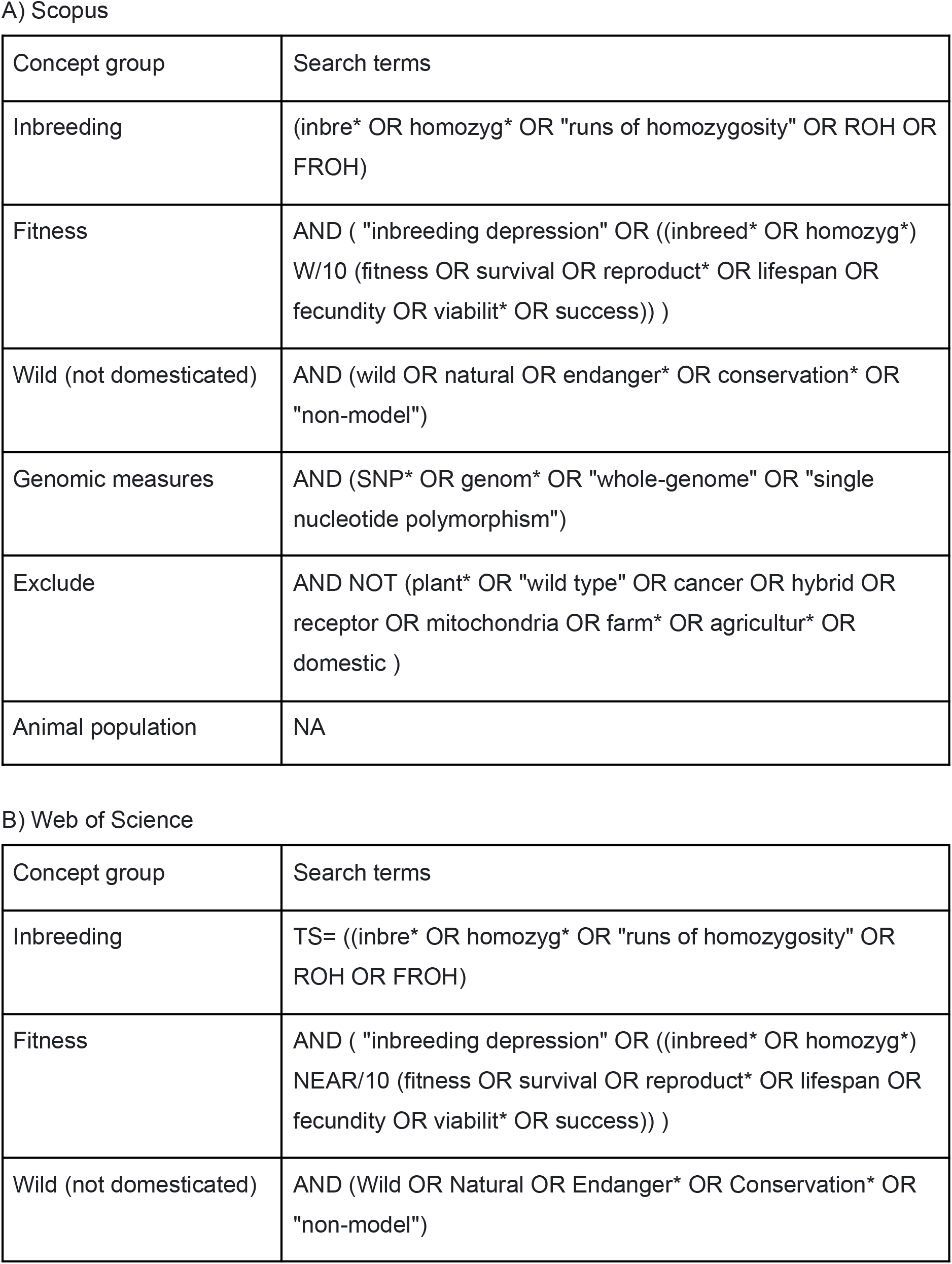
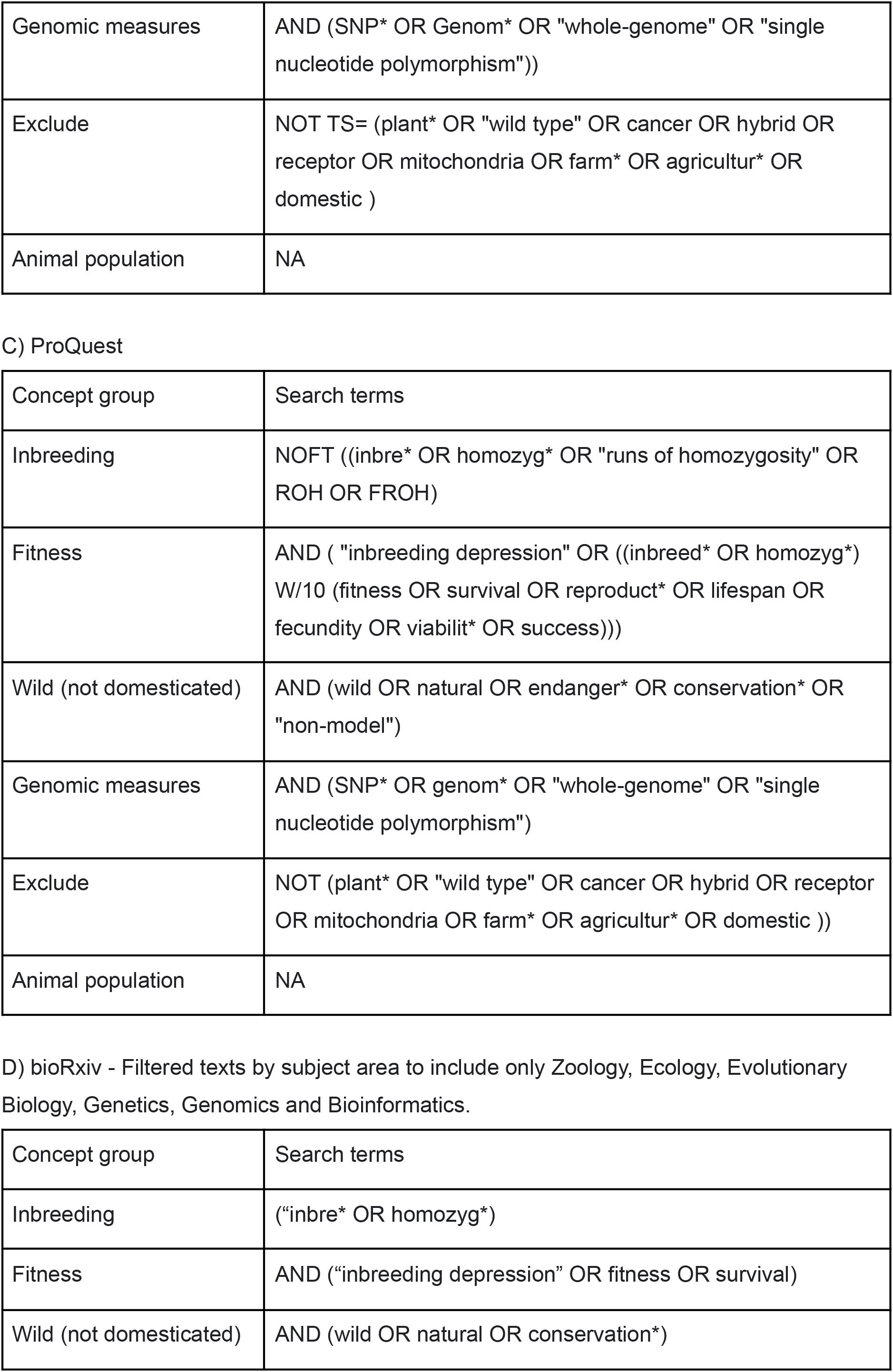
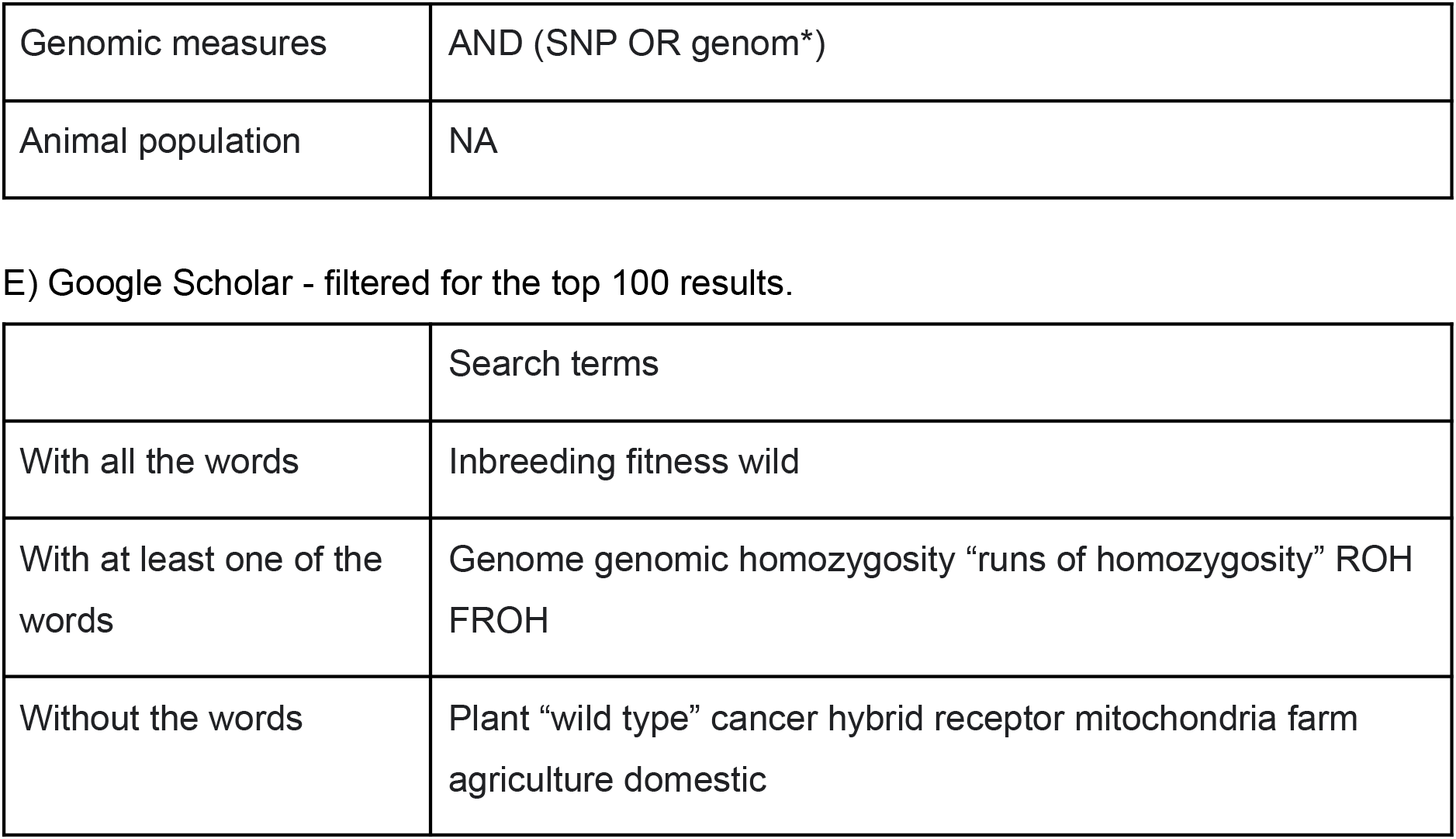
Final Boolean search string used for full literature search in (A) Scopus, (B) Web of Science, (C) ProQuest, (D) bioRxiv and (E) Google Scholar. Each search was limited to texts from 1 Jan 2013 to 17 Jun 2025.

| Concept group | Search terms |
| --- | --- |
| Inbreeding | (inbre* OR homozyg* OR "runs of homozygosity" OR ROH OR FROH) |
| Fitness | AND ( "inbreeding depression" OR ((inbreed* OR homozyg*) W/10 (fitness OR survival OR reproduct* OR lifespan OR fecundity OR viabilit* OR success)) ) |
| Wild (not domesticated) | AND (wild OR natural OR endanger* OR conservation* OR "non-model") |
| Genomic measures | AND (SNP* OR genom* OR "whole-genome" OR "single nucleotide polymorphism") |
| Exclude | AND NOT (plant* OR "wild type" OR cancer OR hybrid OR receptor OR mitochondria OR farm* OR agricultur* OR domestic ) |
| Animal population | NA |

B) Web of Science
| Concept group | Search terms |
| --- | --- |
| Inbreeding | TS= ((inbre* OR homozyg* OR "runs of homozygosity" OR ROH OR FROH) |
| Fitness | AND ( "inbreeding depression" OR ((inbreed* OR homozyg*) NEAR/10 (fitness OR survival OR reproduct* OR lifespan OR fecundity OR viabilit* OR success)) ) |
| Wild (not domesticated) | AND (Wild OR Natural OR Endanger* OR Conservation* OR "non-model") |
| Genomic measures | AND (SNP* OR Genom* OR "whole-genome" OR "single nucleotide polymorphism")) |
| Exclude | NOT TS= (plant* OR "wild type" OR cancer OR hybrid OR receptor OR mitochondria OR farm* OR agricultur* OR domestic ) |
| Animal population | NA |

C) ProQuest
| Concept group | Search terms |
| --- | --- |
| Inbreeding | NOFT ((inbre* OR homozyg* OR "runs of homozygosity" OR ROH OR FROH) |
| Fitness | AND ( "inbreeding depression" OR ((inbreed* OR homozyg*) W/10 (fitness OR survival OR reproduct* OR lifespan OR fecundity OR viabilit* OR success))) |
| Wild (not domesticated) | AND (wild OR natural OR endanger* OR conservation* OR "non-model") |
| Genomic measures | AND (SNP* OR genom* OR "whole-genome" OR "single nucleotide polymorphism") |
| Exclude | NOT (plant* OR "wild type" OR cancer OR hybrid OR receptor OR mitochondria OR farm* OR agricultur* OR domestic )) |
| Animal population | NA |

D) bioRxiv - Filtered texts by subject area to include only Zoology, Ecology, Evolutionary Biology, Genetics, Genomics and Bioinformatics.
| Concept group | Search terms |
| --- | --- |
| Inbreeding | ("inbre* OR homozyg*) |
| Fitness | AND ("inbreeding depression" OR fitness OR survival) |
| Wild (not domesticated) | AND (wild OR natural OR conservation*) |
| Genomic measures | AND (SNP OR genom*) |
| Animal population | NA |
| E) Google Scholar - filtered for the top 100 results. |  |
|  | Search terms |
| With all the words | Inbreeding fitness wild |
| With at least one of the words | Genome genomic homozygosity "runs of homozygosity" ROH<br>FROH |
| Without the words | Plant "wild type" cancer hybrid receptor mitochondria farm<br>agriculture domestic |

**Table S4.2.** Flowchart of questions used for initial screen of title, abstract, and key-words. Studies were only included if their title, abstract, and key-words returned ‘Yes’ answers to all screening questions. Studies that lacked a formal abstract but had a title or informal abstract suggesting at least four of the below questions were answered with a ‘Yes’.

| Initial screen question | Additional notes |
| --- | --- |
| Is there suggestion an animal population was studied? | • If a Linnaean name is given, Google search the taxonomy of the species in question.<br>• Exclude if study is on *Homo sapiens*. |
| Is there suggestion a wild population was studied? | • Include a reference to “feral”.<br>• Include individuals and descendents from translocations, as long as they are freely breeding and in their assumed natural habitat.<br>• Exclude if non-freely breeding.<br>• Exclude if reference to “indigenous”, “commercial”, “domestic”, “zoo” or “captive”. |
| Is there any suggestion that inbreeding was studied? | • Exclude if no specific mention of “inbreeding”. |
| Is there any suggestion that an effect of inbreeding on fitness was studied? | • Only include if explicit mention of traits relating to survival or reproduction OR if it measures fitness but doesn’t state any other exact measures.<br>• Exclude if fitness-related traits are mentioned but not survival or reproduction<br>• Exclude if mutational load alone is used as a proxy for fitness, but include if used in conjunction with other fitness traits. |
| Is there any suggestion that genomic methods using thousands of markers were used to study inbreeding? | • Include by suggestion of the word “genomic” being used: SNPs, or SNP array, or SNP-chip or whole-genome or genome-wide or multi.<br>• Exclude if only “microsatellite markers” are used, however include if used in conjunction with genomic markers.<br>• Exclude if less than 1000 markers are used. |

**Table S4.3.** The 27 studies identified from our systematic review.

| ID | Author & Year | Common name | Linnean name | Taxon | IUCN status | Historical isolation | Inbreeding coefficient | n markers |
| --- | --- | --- | --- | --- | --- | --- | --- | --- |
| 1 | Zilko et al. 2020 | Leadbeater's possum | *Gymnobelideus leadbeateri* | Mammal | CR | Isolated | $F_{GRM}$ | 3,378 |
| 2 | Niskanen et al. 2020 | House sparrow | *Passer domesticus* | Bird | LC | Non-isolated | $F_{ROH}$ | 118,810 |
| 3 | Crossman et al. 2024 | North Atlantic right whale | *Eubalaena glacialis* | Mammal | CR | Non-isolated | $sMLH$ | 26,011 |
| 4 | Troianou et al. 2018 | Red deer | *Cervus elaphus* | Mammal | LC | Non-isolated | $F_{GRM}$ | 37,410 |
| 5 | Hewett et al. 2025 | Red deer | *Cervus elaphus* | Mammal | LC | Non-isolated | $F_{ROH}$ | 37,396 |
| 6 | Stoffel et al. 2021 | Soay sheep | *Ovis aries* | Mammal | LC | Isolated | $F_{ROH}$ | 417,373 |
| 7 | Berenos et al. 2016 | Soay sheep | *Ovis aries* | Mammal | LC | Isolated | $F_{ROH}$ | 13,370 |
| 8 | Hasselgren et al. 2021 | Arctic fox | *Vulpes lagopus* | Mammal | EN | Non-isolated | $F_{ROH}$ | 4,579,222 |
| 9 | Chen et al. 2016 | Florida scrub jay | *Aphelocoma coerulescens* | Bird | VU | Isolated | $F_{UNI}$ | 7,834 |
| 10 | Duntsch et al. 2023 | New Zealand stitchbird | *Notiomystis cincta* | Bird | VU | Isolated | $F_{ROH}$ | 41,195 |
| 11 | Foster et al. 2021 | Kakapo | *Strigops habroptila* | Bird | CR | Isolated | $F_{ROH}$ | 12,241 |
| 12 | Hoffman et al. 2014 | Harbor seal | *Phoca vitulina* | Mammal | LC | Non-isolated | $sMLH$ | 14,585 |
| 13 | Huisman et al. 2016 | Red deer | *Cervus elaphus* | Mammal | LC | Non-isolated | $F_{GRM}$ | 37,410 |
| 14 | Pemberton et al. 2017 | Soay sheep | *Ovis aries* | Mammal | LC | Isolated | $F_{GRM}$ | 37,037 |
| 15 | Kardos et al. 2023 | Killer whale | *Orcinus orca* | Mammal | DD | Non-isolated | $F_{ROH}$ | 7,029 |
| 16 | Colpitts et al. 2023 | Feral horses | *Equus caballus* | Mammal | LC | Non-isolated | $F_{ROH}$ | 30,415 |
| 17 | Clark et al. 2024 | Eastern massasauga | *Sistrurus catenatus* | Reptile | LC | Non-isolated | $F_{GRM}$ | 2,176 |
| 18 | Kubacka et al. 2025 | Aquatic warbler | *Acrocephalus paludicola* | Bird | VU | Non-isolated | $F_{ROH}$ | 2,948 |
| 19 | Harrison et al. 2019 | Helmeted honeyeater | *Lichenostomus melanops* | Bird | LC | Isolated | $F_{HOM}$ | 11,059 |
| 20 | Paijmans et al. 2024 | Antarctic fur seals | *Arctocephalus gazella* | Mammal | LC | Non-isolated | $F_{ROH}$ | 75,101 |
| 21 | Stoffel et al. 2021 | Soay sheep | *Ovis aries* | Mammal | LC | Isolated | $F_{ROH}$ | 430,702 |
| 22 | Afonso et al. 2023 | Bermuda petrel | *Pterodroma cahow* | Bird | EN | Isolated | $sMLH$ | 12,484 |
| 23 | Hasik et al. 2025 | Red deer | *Cervus elaphus* | Mammal | LC | Non-isolated | $F_{ROH}$ | 37,396 |
| 24 | Vonholdt et al. 2023 | Grey wolf | *Canis lupus* | Mammal | LC | Non-isolated | $F_{ROH}$ | 24,235 |
| 25 | Yates et al. 2019 | Brook trout | *Salvelinus fontinalis* | Fish | LC | Non-isolated | $H_O$ | 5,858 |
| 26 | Hasselgren et al. 2024 | Arctic fox | *Vulpes lagopus* | Mammal | EN | Non-isolated | $H_O$ | 10,880,359 |
| 27 | Lee et al. 2026 | Seychelles warbler | *Acrocephalus sechellensis* | Bird | VU | Isolated | $F_{ROH}$ | 1,890,803 |

## References

1. Ablondi, M., Summer, A., Stocco, G., Finocchiaro, R., van Kaam, J.-T., Cassandro, M., et al. (2023). The role of inbreeding depression on productive performance in the Italian Holstein breed. J Anim Sci, 101, skad382.

2. Alemu, S.W., Kadri, N.K., Harland, C., Faux, P., Charlier, C., Caballero, A., et al. (2020). An evaluation of inbreeding measures using a whole-genome sequenced cattle pedigree. Heredity 2020 126:3, 126, 410–423.

3. Altman, D.G. & Bland, J.M. (2011). How to obtain the confidence interval from a P value. BMJ, 343, d2090.

4. Auclair, L., Vanpé, C., Chapron, G., Quenette, P.-Y. & Robert, A. (2025). Inbreeding Depression Across Multiple Life-History Traits in a Long-Lived Mammal. Molecular Ecology, 34, e70123.

5. Balloux, F., Amos, W. & Coulson, T. (2004). Does heterozygosity estimate inbreeding in real populations? Molecular Ecology, 13, 3021–3031.

6. Beccardi, M., Pen, I., Bichet, C., Tschirren, B. & Vedder, O. (2024). Inbreeding accelerates reproductive senescence, but not survival senescence, in a precocial bird. Journal of Animal Ecology, 93, 1972–1982.

7. Bonnet, T., Morrissey, M.B., de Villemereuil, P., Alberts, S.C., Arcese, P., Bailey, L.D., et al. (2022). Genetic variance in fitness indicates rapid contemporary adaptive evolution in wild animals. Science, 376, 1012–1016.

8. Borenstein, M., Hedges, L.V., Higgins, J.P.T. & Rothstein, H.R. (2009). Introduction to Meta Analysis. Wiley.

9. Caballero, A., Fernández, A., Villanueva, B. & Toro, M.A. (2022). A comparison of marker-based estimators of inbreeding and inbreeding depression. Genet Sel Evol, 54, 82.

10. Caballero, A., Villanueva, B. & Druet, T. (2021). On the estimation of inbreeding depression using different measures of inbreeding from molecular markers. Evolutionary Applications, 14, 416–428.

11. Chan, Y.F. (2022). The impacts of environmental conditions on inbreeding depression: a meta-analysis. The University of Liverpool.

12. Chapman, J.R., Nakagawa, S., Coltman, D.W., Slate, J. & Sheldon, B.C. (2009). A quantitative review of heterozygosity–fitness correlations in animal populations. Molecular Ecology, 18, 2746–2765.

13. Charlesworth, B. & Hughes, K.A. (1996). Age-specific inbreeding depression and components of genetic variance in relation to the evolution of senescence. Proceedings of the National Academy of Sciences, 93, 6140–6145.

14. Charlesworth, D. & Willis, J.H. (2009). The genetics of inbreeding depression. Nature Reviews Genetics, 10, 783–796.

15. Clark, M.I., Hileman, E.T., Moore, J.A., Faust, L.J., Junge, R.E., Reid, B.N., et al. (2025). Inbreeding reduces fitness in spatially structured populations of a threatened rattlesnake. Proceedings of the National Academy of Sciences, 122, e2501745122.

16. Cohen, J. (1960). A Coefficient of Agreement for Nominal Scales. Educational and Psychological Measurement, 20, 37–46.

17. Crossman, C.A., Hamilton, P.K., Brown, M.W., Conger, L.A., George, R.C., Jackson, K.A., et al. (2024). Effects of inbreeding on reproductive success in endangered North Atlantic right whales. R Soc Open Sci., 11, 240490.

18. Culina, A., Adriaensen, F., Bailey, L.D., Burgess, M.D., Charmantier, A., Cole, E.F., et al. (2021). Connecting the data landscape of long-term ecological studies: The SPI-Birds data hub. Journal of Animal Ecology, 90, 2147–2160.

19. Duntsch, L., Whibley, A., Brekke, P., Ewen, J.G. & Santure, A.W. (2021). Genomic data of different resolutions reveal consistent inbreeding estimates but contrasting homozygosity landscapes for the threatened Aotearoa New Zealand hihi. Molecular Ecology, 30, 6006–6020.

20. Duntsch, L., Whibley, A., de Villemereuil, P., Brekke, P., Bailey, S., Ewen, J.G., et al. (2023). Genomic signatures of inbreeding depression for a threatened Aotearoa New Zealand passerine. Mol Ecol, 32, 1893–1907.

21. Dussex, N., van der Valk, T., Morales, H.E., Wheat, C.W., Díez-del-Molino, D., von Seth, J., et al. (2021). Population genomics of the critically endangered kākāpō. Cell Genomics, 1, 100002.

22. Ebel, E.R. & Phillips, P.C. (2016). Intrinsic differences between males and females determine sex-specific consequences of inbreeding. BMC Evol Biol, 16, 36.

23. Enders, L.S. & Nunney, L. (2010). Sex-specific effects of inbreeding in wild-caught *Drosophila melanogaster* under benign and stressful conditions. Journal of Evolutionary Biology, 23, 2309–2323.

24. Forutan, M., Ansari Mahyari, S., Baes, C., Melzer, N., Schenkel, F.S. & Sargolzaei, M. (2018). Inbreeding and runs of homozygosity before and after genomic selection in North American Holstein cattle. BMC Genomics, 19, 98.

25. Foster, Y., Dutoit, L., Grosser, S., Dussex, N., Foster, B.J., Dodds, K.G., et al. (2021). Genomic signatures of inbreeding in a critically endangered parrot, the kākāpō. G3: Genes, Genomes, Genetics, 11.

26. Grieshop, K., Maurizio, P.L., Arnqvist, G. & Berger, D. (2021). Selection in males purges the mutation load on female fitness. Evolution Letters, 5, 328–343.

27. Haddaway, N.R., Page, M.J., Pritchard, C.C. & McGuinness, L.A. (2022). PRISMA2020: An R package and Shiny app for producing PRISMA 2020-compliant flow diagrams, with interactivity for optimised digital transparency and Open Synthesis. Campbell Systematic Reviews, 18, e1230.

28. Hasik, A.Z., Hewett, A.M., Maris, K., Morris, S.J., Morris, A., Albery, G.F., et al. (2025). Parasite-mediated inbreeding depression in wild red deer. Heredity, 134, 637–644.

29. Hasselgren, M., Angerbjörn, A., Eide, N.E., Erlandsson, R., Flagstad, Ø., Landa, A., et al. (2018). Genetic rescue in an inbred Arctic fox (*Vulpes lagopus*) population. Proceedings of the Royal Society B: Biological Sciences, 285, 20172814.

30. Hewett, A.M., Johnston, S.E., Albery, G.F., Morris, A., Morris, S.J. & Pemberton, J.M. (2025). Fine-scale spatial variation in fitness, inbreeding, and inbreeding depression in a wild ungulate. Evol Lett, 9, 292–301.

31. Huisman, J., Kruuk, L.E.B., Ellisa, P.A., Clutton-Brock, T. & Pemberton, J.M. (2016). Inbreeding depression across the lifespan in a wild mammal population. Proceedings of the National Academy of Sciences of the United States of America, 113, 3585–3590.

32. Janicke, T., Vellnow, N., Sarda, V. & David, P. (2013). Sex-specific inbreeding depression depends on the strength of male–male competition. Evol, 67, 2861–2875.

33. Kardos, M., Taylor, H.R., Ellegren, H., Luikart, G. & Allendorf, F.W. (2016). Genomics advances the study of inbreeding depression in the wild. Evolutionary Applications, 9, 1205–1218.

34. Keller, L.F. & Waller, D.M. (2002). Inbreeding effects in wild populations. Trends in Ecology and Evolution, 17, 230–241.

35. Kyriazis, C.C., Robinson, J.A. & Lohmueller, K.E. (2025). Long runs of homozygosity are reliable genomic markers of inbreeding depression. Trends in Ecology & Evolution, 40, 874–884.

36. Lande, R. (1988). Genetics and Demography in Biological Conservation. Science, 241, 1455–1460.

37. Landis, J. & Koch, G. (1977). The measurement of observer agreement for categorical data. Biometrics, 33, 159–174.

38. Lavanchy, E. & Goudet, J. (2023). Effect of reduced genomic representation on using runs of homozygosity for inbreeding characterization. Molecular Ecology Resources, 23, 787–802.

39. Lavanchy, E., Weir, B.S. & Goudet, J. (2024). Detecting inbreeding depression in structured populations. Proc Natl Acad Sci U S A, 121, e2315780121.

40. Lee, K.G.L., Pinto, A., Dong, S., González-Mollinedo, S., Lee, C.Z., Slate, J., et al. (2026). Inbreeding depression by polygenic load following a severe population bottleneck.

41. Mallet, M.A. & Chippindale, A.K. (2011). Inbreeding reveals stronger net selection on *Drosophila melanogaster* males: implications for mutation load and the fitness of sexual females. Heredity, 106, 994–1002.

42. Martikainen, K., Sironen, A. & Uimari, P. (2018). Estimation of intrachromosomal inbreeding depression on female fertility using runs of homozygosity in Finnish Ayrshire cattle. Journal of Dairy Science, 101, 11097–11107.

43. Michonneau, F., Brown, J.W. & Winter, D.J. (2016). rotl: an R package to interact with the Open Tree of Life data. Methods in Ecology and Evolution, 7, 1476–1481.

44. Moher, D., Liberati, A., Tetzlaff, J., Altman, D.G. & Group, T.P. (2009). Preferred Reporting Items for Systematic Reviews and Meta-Analyses: The PRISMA Statement. PLOS Medicine, 6, e1000097.

45. Mota, L.F.M., Carvajal, A.B., Silva Neto, J.B., Díaz, C., Carabaño, M.J., Baldi, F., et al. (2024). Assessment of inbreeding coefficients and inbreeding depression on complex traits from genomic and pedigree data in Nelore cattle. BMC Genomics, 25, 944.

46. Nakagawa, S. & Cuthill, I.C. (2007). Effect size, confidence interval and statistical significance: a practical guide for biologists. Biological Reviews, 82, 591–605.

47. Nakagawa, S., Lagisz, M., O’Dea, R.E., Rutkowska, J., Yang, Y., Noble, D.W.A., et al. (2021). The orchard plot: Cultivating a forest plot for use in ecology, evolution, and beyond. In: Research Synthesis Methods. John Wiley and Sons Ltd, pp. 4–12.

48. Nakagawa, S. & Schielzeth, H. (2013). A general and simple method for obtaining R 2 from generalized linear mixed-effects models. Methods in Ecology and Evolution, 4, 133–142.

49. Nietlisbach, P., Muff, S., Reid, J.M., Whitlock, M.C. & Keller, L.F. (2019). Nonequivalent lethal equivalents: Models and inbreeding metrics for unbiased estimation of inbreeding load. Evolutionary Applications, 12, 266–279.

50. Nussey, D.H., Pemberton, J., Donald, A. & Kruuk, L.E.B. (2006). Genetic consequences of human management in an introduced island population of red deer (*Cervus elaphus*). Heredity, 97, 56–65.

51. Ouzzani, M., Hammady, H., Fedorowicz, Z. & Elmagarmid, A. (2016). Rayyan—a web and mobile app for systematic reviews. Systematic Reviews 2016 5:1, 5, 1–10.

52. Paradis, E. & Schliep, K. (2019). ape 5.0: an environment for modern phylogenetics and evolutionary analyses in R. Bioinformatics, 35, 526–528.

53. Peripolli, E., Munari, D.P., Silva, M.V.G.B., Lima, A.L.F., Irgang, R. & Baldi, F. (2017). Runs of homozygosity: current knowledge and applications in livestock. Animal Genetics, 48, 255–271.

54. Purcell, S., Neale, B., Todd-Brown, K., Thomas, L., Ferreira, M.A.R., Bender, D., et al. (2007). PLINK: a tool set for whole-genome association and population-based linkage analyses. Am J Hum Genet, 81, 559–575.

55. Rosenberg, M.S., Rothstein, H.R. & Gurevitch, J. (2013). Effect Sizes: Conventional Choices and Calculations. In: Handbook of Meta-analysis in Ecology and Evolution (eds. Koricheva, J., Gurevitch, J. & Mengersen, K.). Princeton University Press, pp. 61–71.

56. Slate, J., David, P., Dodds, K.G., Veenvliet, B.A., Glass, B.C., Broad, T.E., et al. (2004). Understanding the relationship between the inbreeding coefficient and multilocus heterozygosity: Theoretical expectations and empirical data. Heredity, 93, 255–265.

57. Stoffel, M.A., Esser, M., Kardos, M., Humble, E., Nichols, H., David, P., et al. (2016). inbreedR: an R package for the analysis of inbreeding based on genetic markers. Methods in Ecology and Evolution, 7, 1331–1339.

58. Stoffel, M.A., Johnston, S.E., Pilkington, J.G. & Pemberton, J.M. (2021). Genetic architecture and lifetime dynamics of inbreeding depression in a wild mammal. Nature Communications, 12, 1–10.

59. Trask, A.E., Ferrie, G.M., Wang, J., Newland, S., Canessa, S., Moehrenschlager, A., et al. (2021). Multiple life-stage inbreeding depression impacts demography and extinction risk in an extinct-in-the-wild species. Sci Rep, 11, 682.

60. Tsujimoto, D., Takayanagi, M., Ishii, J., Sakashita, R., Horikoshi, K., Suzuki, H., et al. (2025). Genetic purging in an island-endemic pigeon recovering from the brink of extinction. Commun Biol, 8, 1051.

61. VanRaden, P.M. (2008). Efficient methods to compute genomic predictions. Journal of Dairy Science, 91, 4414–4423.

62. Vega-Trejo, R., de Boer, R.A., Fitzpatrick, J.L. & Kotrschal, A. (2022). Sex-specific inbreeding depression: A meta-analysis. Ecology Letters, 25, 1009–1026.

63. Viechtbauer, W. (2010). Conducting Meta-Analyses in R with the metafor Package. Journal of Statistical Software, 36, 1–48.

64. Wang, J. (2016). Pedigrees or markers: Which are better in estimating relatedness and inbreeding coefficient? Theor Popul Biol, 107, 4–13.

65. Yang, J., Lee, S.H., Goddard, M.E. & Visscher, P.M. (2011). GCTA: A Tool for Genome-wide Complex Trait Analysis. American Journal of Human Genetics, 88, 76.

66. Yengo, L., Zhu, Z., Wray, N.R., Weir, B.S., Yang, J., Robinson, M.R., et al. (2017). Detection and quantification of inbreeding depression for complex traits from SNP data. Proc Natl Acad Sci U S A, 114, 8602–8607.

